# Syntrophic co-culture amplification of production phenotype for high-throughput screening of microbial strain libraries

**DOI:** 10.1101/518639

**Authors:** Tatyana E. Saleski, Alissa R. Kerner, Meng Ting Chung, Corine M. Jackman, Azzaya Khasbaatar, Katsuo Kurabayashi, Xiaoxia Nina Lin

**Affiliations:** Department of Chemical Engineering, University of Michigan, Ann Arbor, MI, USA; Department of Mechanical Engineering, University of Michigan, Ann Arbor, MI, USA; Department of Electrical Engineering and Computer Science, University of Michigan, Ann Arbor, MI, USA; Department of Biomedical Engineering, University of Michigan, Ann Arbor, MI, USA

**Author notes:** **Correspondence:** Xiaoxia Nina Lin, G054W NCRC Bldg 28, 2800 Plymouth Rd., Ann Arbor, MI 48109-2800.

**Keywords:** High-throughput strain screening, cross-feeding, microbial co-culture, droplet microfluidics, biosensor, signal amplification, industrial microbiology, metabolic engineering

## Abstract

Microbes can be engineered to synthesize a wide array of bioproducts, yet production phenotype evaluation remains a frequent bottleneck in the design-build-test cycle where strain development requires iterative rounds of library construction and testing. Here, we present <u>S</u>y<u>n</u>tr<u>o</u>phic <u>C</u>o-culture <u>A</u>mplification of <u>P</u>roduction phenotype (SnoCAP). Through a metabolic cross-feeding circuit, the production level of a target molecule is translated into highly distinguishable co-culture growth characteristics, which amplifies differences in production into highly distinguishable growth phenotypes. We demonstrate SnoCAP with the screening of *Escherichia coli* strains for production of two target molecules: 2-ketoisovalerate, a precursor of the drop-in biofuel isobutanol, and L-tryptophan. The dynamic range of the screening can be tuned by employing an inhibitory analog of the target molecule. Screening based on this framework requires compartmentalization of individual producers with the sensor strain. We explore three formats of implementation with increasing throughput capability: confinement in microtiter plates (10^2^-10^4^ assays/experiment), spatial separation on agar plates (10^4^-10^5^ assays/experiment), and encapsulation in microdroplets (10^5^-10^7^ assays/experiment). Using SnoCAP, we identified an efficient isobutanol production strain from a random mutagenesis library, reaching a final titer that is 5-fold higher than that of the parent strain. The framework can also be extended to screening for secondary metabolite production using a push-pull strategy. We expect that SnoCAP can be readily adapted to the screening of various microbial species, to improve production of a wide range of target molecules.

**Highlights:**

- A high-throughput screening platform based on cross-feeding auxotrophs was developed.
- Compartmentalization was implemented in three formats: microplates, agar plates, and microdroplets.
- Utility of the screening was demonstrated for two proof-of-concept target molecules: 2-ketoisovalerate and L-tryptophan.
- The assay dynamic range was tuned by addition of an inhibitory analog.
- The screening was applied to identify a strain from a chemically mutagenized library that produces 5-fold higher isobutanol titer than the parent strain.

## 1. Introduction

Advances in genome engineering design and construction technologies have enabled rapid generation of diverse microbial strains that efficiently explore the genotype space (Haimovich, Muir, and Isaacs 2015). Characterizing these strains, however, is often a bottleneck in the design-build-test cycle of synthetic biology. In particular, concerning the development of strains for production of many target molecules, increasingly large and complex strain libraries can be created, yet the throughput of screening for identifying top-performing variants is limited, sometimes lagging several orders of magnitude behind the construction phase. Traditionally, for molecules that lack chromogenic or fluorescent properties, which include an ever-expanding array of pharmaceuticals, specialty and commodity chemicals, and biofuels, metabolic engineers must rely on chromatography or mass spectrometry quantification. Automation can help to increase the throughput of these assays, yet this brute force approach usually poses high capital investment and space requirements. Biosensor-based high-throughput screenings seek to address this challenge by converting target molecule production level into a conspicuous phenotype such as growth or fluorescence, either within the production cell itself or in a partner strain. The latter approach enables the sensing of extracellular secretion levels and reduces the interference between the production and sensing functionalities. The sensing machinery generally consists of proteins, nucleic acid molecules, or whole cells that respond to the target molecule in a dose-dependent manner and produce a detectable read-out (Dietrich, McKee, and Keasling 2010; Park, Tsai, and Chen 2013; Lin, Wagner, and Alper 2017).

One class of whole-cell biosensor consists of auxotrophic strains which are unable to produce an essential metabolite and whose growth characteristics, therefore, change in response to changing concentrations of the target molecule in the surrounding environment. Auxotrophic microbial strains have been identified or constructed and utilized as biosensors for a variety of amino acids (Payne, Bell, and Higgins 1977; Chalova et al. 2007; Chalova et al. 2009; Bertels, Merker, and Kost 2012; Kim et al. 2014), as well as certain vitamins (Beadle and Tatum 1941; Burkholder 1951) and co-factors (Lindner et al. 2018). For these sensor strains, the final cell density reached varies linearly with the starting concentration of the focal molecule, until the concentration of the molecule saturates what is required for attaining the maximum growth and further increase in concentration of the molecule does not lead to further increase in final cell density. Pfleger et al. (2007) applied auxotrophic biosensors to high-throughput screening of production strain libraries by developing a fluorescent mevalonate auxotroph whose growth reports on production strain performance.

Tepper & Shlomi (2011) have established a computational framework to predict gene deletions that can be used to produce auxotrophic strains for use as biosensors. For *Escherichia coli*, for instance, they predict 53 molecules for which auxotrophic biosensor strains could be created. Furthermore, for molecules for which no auxotrophic biosensor is available, they present a strategy to engineer the producer strain by gene knockout so that production of the target molecule is coupled to that of a proxy metabolite, for which a biosensor does exist. It is also possible to construct auxotrophs that depend on molecules that are not normally used for growth by engineering ligand-dependent essential genes (Lopez and Anderson 2015).

Despite the array of auxotrophic biosensors available, a limitation in applying them to high-throughput screening is that they generally have narrow dynamic ranges, confined to low concentrations of the focal molecules. Thus, although these molecules are essential for growth, they do not directly confer a selective advantage if the strains are producing more than small quantities of the molecule. For example, when auxotrophic *E. coli* strains are employed for amino acid quantification, the linear range is typically in the tens of micromolar. This small dynamic range limitation can sometimes be overcome by dilution of the samples (Pfleger et al. 2007; Niu et al. 2011), but this makes screening more cumbersome, lowering throughput.

Syntrophic co-cultures, consisting of auxotrophic strains in a cross-feeding circuit that enables co-growth, have been used historically by microbiologists as a tool to interrogate biochemical pathways (Winkler, van Doorn, and Royers 1952; Nurmikko 1956) and for assessment of whether a certain metabolite is produced by a strain of interest (Fildes 1956). Here, we describe <u>S</u>yntr<u>o</u>phic <u>C</u>o-culture <u>A</u>mplification of <u>P</u>roduction phenotype (SnoCAP) for high-throughput screening of production strains via colocalization with a partner strain (Fig. 1). One strain, the “sensor”, is auxotrophic for the target molecule; its ability to grow depends upon the amount of target molecule excreted by the other strain, the “secretor.” The secretor is auxotrophic for an orthogonal molecule supplied by the sensor strain. In model microbial systems, it has been shown that changes in the secretion or uptake characteristics of either partner of a cross-feeding pair determine the resulting composition of the co-culture as well as its overall co-growth rate (Kerner et al. 2012; Zhang and Reed 2014). We predict that a secretor strain with improved production rate will lead to faster growth and an increased final sensor-to-secretor ratio (Fig. 1A; Section 3, Calculation). We demonstrate the utility of the co-culture screening strategy for high-throughput screening of *E. coli* strain libraries and explore three methods for compartmentalization of unique secretor strains with the sensor strain: confinement in wells of microtiter plates, spatial separation as colonies on agar plates, and encapsulation in water-in-oil microdroplets (Fig. 1B).

**Figure 1:**
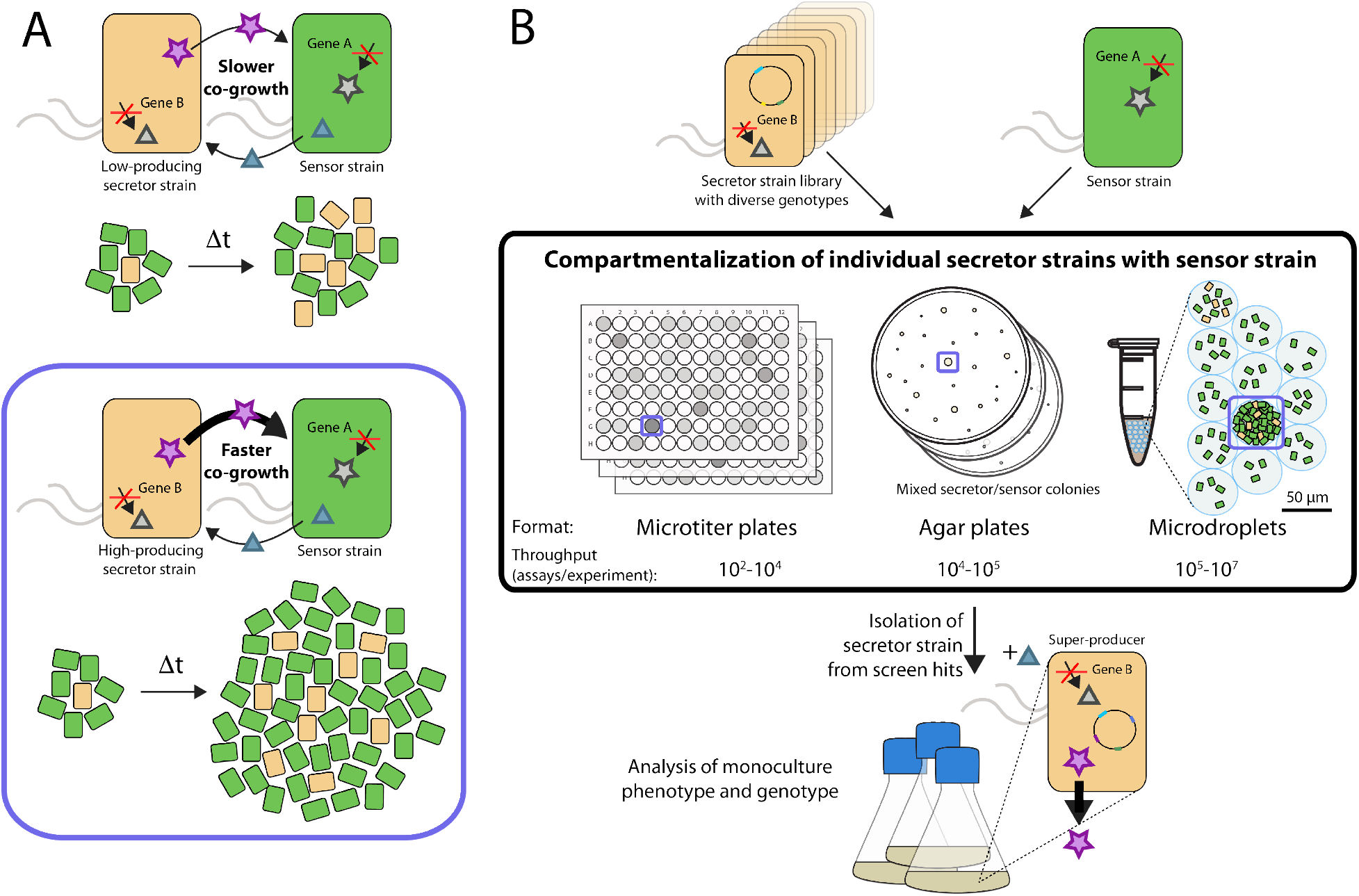
Overall schematic of screening strategy. (A) Production improvement leads to increased co-growth and increased final sensor-to-secretor ratio. Δt represents a change in time. The purple star represents the molecule that one wishes to overproduce. The blue triangle represents a secondary cross-fed metabolite. (B) Screening implementation formats explored in this study: separating secretor cells of unique genotype by compartmentalization in wells of microplates, spatial separation on agar plates, or confinement in microdroplets. After growth, secretor strains are isolated from well-grown co-cultures for further analysis in monocultures.

We expect the cross-feeding system to have two major advantages for high-throughput screening over traditional auxotrophic biosensor implementation: i) amplification of production improvement through the co-culture’s exponential growth, and ii) a wider and more tunable dynamic range. The former lies in the co-culture’s amplification of differences between library members (Section 3, Calculation). Since strain development generally proceeds through incremental improvements, it is critical to be able to discern modest increases in production. Amplification of differences through the cross-feeding metabolic circuit makes relatively small improvements conspicuous. The traditional auxotrophic biosensor approach has a linear change in sensor output (optical density or fluorescence) in response to changing target molecule concentration, providing no amplification. The second advantage is achieved because, when neither of the cross-fed molecules is exogenously supplied, the production rates of the cross-fed molecules generally limit total co-culture growth, keeping the target molecule within the dynamic range of the sensor strain. This is in contrast to direct application of the sensor to the supernatant of a secretor strain when the level of target molecule often exceeds sensor dynamic range, necessitating cumbersome strategies such as dilution of sample supernatant.

Screening by SnoCAP requires the producer to be auxotrophic for a molecule that the sensor strain can supply. Unless the strain is naturally auxotrophic for an appropriate molecule, the parent secretor needs to be genetically modified to effect the auxotrophy. This can be achieved by targeted gene deletion, as shown in this work, or by mutagenesis and screening. It could also involve addition of genes that enable utilization of the cross-fed molecule. Following the library construction and SnoCAP screening, the auxotrophy can be easily reversed, if needed, by targeted reintroduction of the previously deleted gene or by phage transduction from the wildtype strain.

## 2. Materials and Methods

Key elements are described here. Additional details are provided in the Supplementary Information.

### 2.1 Strains and plasmids

Strains and plasmids used are listed in Table S1. JCL16, JCL260, pSA65 and pSA69 (Atsumi, Hanai, and Liao 2008; Atsumi et al. 2010) were provided by James Liao, UCLA. Keio strains were obtained from the *E. coli* genetic stock center (CGSC, http://cgsc2.biology.yale.edu/). The pTGD plasmid (Tyo, Ajikumar, and Stephanopoulos 2009) was provided by Keith Tyo, Northwestern University. pET-ara-mCherry was constructed by Jihyang Park (Park et al. 2011) and pSAS31 by Scott Scholz (Scholz et al. 2018). The Δ*intC*::*yfp-cat* construct was originally obtained from the DS1-Y strain, from the Balaban group, Hebrew University of Israel. pBT1-proD-mCherry was a gift from Michael Lynch (Addgene plasmid #65823).

### 2.2 Cell preparation for cross-feeding screening

Strains were maintained as glycerol stocks at −80 C and streaked from cryostocks on LB agar plates with the appropriate antibiotics. Colonies were then picked into liquid LB Lennox with appropriate antibiotics and grown to stationary phase (16-18 h, 37 °C, 250 rpm). For microtiter plate and agar plate assays, the stationary phase cells were harvested in 1 mL aliquots by centrifugation at 12,000 g for 1 min. The cells were washed twice with 1 mL of 1x M9 salts without amino acids, and resuspended to an optical density corresponding to ~10^9^ CFU/mL based on OD_600_ measurement in a VersaMax microplate reader (Molecular Devices). For the droplet assay, the stationary phase cultures were subcultured in 10 mL culture volumes into exponential phase (~3.5 h for JCL260-based strains and ~2 h for K12-based strains). The cells were harvested by centrifugation at 5,000 g for 5 min, washed twice in 1x M9 salts, and resuspended in the culturing medium (M9IPG with norvaline at the specified concentration). Cell density was then determined based on an OD_600_ to CFU calibration. Cell inocula were kept at room temperature and each culture was combined just before droplet generation. Each inoculum culture was serially diluted and spot plated on LB plates with kanamycin to verify CFU concentration.

### 2.3 Media

M9IPG, consisting of M9 salts (47.8 mM Na_2_HPO_4_, 22.0 mM KH_2_PO_4_, 8.55 mM NaCl, 9.35 mM NH_4_Cl, 1 mM MgSO_4_, 0.3 mM CaCl_2_), micronutrients (2.91 nM (NH_4_)_2_MoO_4_, 401.1 nM H_3_BO_3_, 30.3 nM CoCl_2_, 9.61 nM CuSO_4_, 51.4 nM MnCl_2_, 6.1 nM ZnSO_4_, 0.01 mM FeSO_4_), thiamine HCl (3.32 μM) and dextrose (D-glucose) at the stated concentrations, was used as the base medium. When specified, 5 g/L yeast extract was added to the medium. Antibiotics were used at the following concentrations: ampicillin, 100 μg/mL; kanamycin, 50 μg/mL; tetracycline, 10 μg/mL; chloramphenicol, 20 or 80 μg/mL. All amino acids were the enantiopure L-isomer, except for norvaline, which was a racemic form (Thermo Fisher Scientific).

### 2.4 Microplate assay

M9IPG with 20 g/L glucose, 3 mM isoleucine, and 50 μg/mL kanamycin was used for all microplates in the 2-KIV screen. No other antibiotics were added. When noted, the medium contained IPTG at 0.1 mM concentration. M9IPG with 4 g/L glucose and no antibiotics were used for all microplates in the tryptophan screening system. When noted, cultures were supplemented with the stated concentrations of tryptophan and histidine. Cells were prepared as described above and then inoculated into medium. For monocultures, the cells were inoculated 1:100 by volume unless otherwise specified. For co-cultures, each strain was inoculated 1:200 by volume. Cultures were vortexed and then distributed into a 96-well clear microplate (Brand), 200 μL per well. Microplate lids were coated with a solution of 0.5% Triton X-100 in 20% ethanol to reduce condensation and lids were fastened with tape. Microplates were incubated at 37 °C, with shaking in a VersaMax plate reader (Molecular Devices), with absorbance readings at 600 nm taken every 10 min. μ_max_ was calculated via linear regression of natural log of OD_600_ values (after subtracting blank values) vs. time; regression was performed over the time intervals corresponding to early exponential growth phase.

### 2.5 Agar plate assay

#### 2.5.1 Plate setup

Plates were either 10 cm round Petri dishes, or 24.5 cm square bioassay dishes. M9IPG with 5 g/L glucose, 3 mM isoleucine, 12 g/L agar and 50 μg/mL kanamycin was used for all plates in the 2-KIV screen. M9IPG with 4 g/L glucose, 12 g/L agar and no antibiotics was used for all microplates in the tryptophan screen. The prepared sensor cells were diluted 10^−1^, secretor cells were serially diluted to the desired concentrations. On round plates, 100 μL of 10^−1^ diluted sensor cells and 200 μL 10^−6^ diluted secretor cells were spread with glass beads. On square plates, 1 mL of 10^−1^ diluted sensor cells and a combination of secretor cells totaling ~10^4^ CFU were spread with glass beads. Plates were allowed to dry thoroughly before incubation to ensure good separation of colonies. 10^−6^ dilutions of secretor strains were also plated on LB plates to determine LB-CFU counts. The LB-CFU counts were used to determine model library percentages.

#### 2.5.2 Scanning and analysis

Automated scanning and ScanLag analysis was performed according to the method of (Levin-Reisman et al. 2010; Levin-Reisman, Fridman, and Balaban 2014), using Epson Perfection V37 photo scanners. Custom holders were 3D-printed so that plates would be located in consistent locations between experiments. Plates were covered with sterile black felt prior to incubation and scanning. Scanners and plates were incubated at 35 °C to accommodate recommended scanner operating temperature. Images were taken every 30 min. for the duration of the incubation period. Following colony growth, images were aligned and colonies detected. Colonies that had merged by the end of the culture period were eliminated from the growth profile plots.

### 2.6 Microdroplet Assay

#### 2.6.1 Encapsulation and cultivation

Cells were prepared as described above. Individual strain inocula were kept at room temperature (storage on ice decreases viability for co-culture growth) and were combined with each other immediately before each set of encapsulation. The strains were combined with additional medium to achieve a total secretor cell loading of λ_Secretor_ = 0.1 cell per droplet and sensor cell loading of λ_Sensor_ = 5 cells per droplet for 55 μm diameter droplets, or λ_Sensor_ = 15 cells per droplet for ~125 μm diameter droplets. M9IPG medium with 20 g/L glucose was supplemented with 50 μg/mL kanamycin, 0.1 mM IPTG, and norvaline at the specified concentration. The inoculated cell culture and Novec HFE-7500 fluorinated oil (3 M) with 2% (w/w) PEG-PFPE amphiphilic block copolymer surfactant (Ran Biotechnologies, 008-FluoroSurfactant) were loaded into separate syringes (BD, 1 mL and 5 mL, respectively) and injected with 23 gauge needles through PTFE tubing (0.022” ID, Cole-Parmer) into a flow-focusing droplet generation device (Fig. S9A) using syringe pumps (Kent Scientific) with flow rates of 10 μL/min and 45 μL/min, respectively. With a device with channel height of 50 μm and aqueous and oil channel widths of 25 μm, these flow rates produced droplets with diameter of ~55 μm. The emulsion was collected in 1.5 mL Eppendorf tubes in 10 min aliquots (~150 μL of emulsion). Excess oil was removed, except for ~100 μL, and the capped Eppendorf tubes were then incubated at 37 °C. Overall population-level growth characteristics were monitored by incubating 100 μL of droplets with 50 μL additional fluorinated oil with surfactant in a black clear-bottom microplate (Greiner), sealed with a Mylar plate sealer (Thermo Scientific), in a microplate reader (BioTek Synergy H1) at 37 °C, reading fluorescence every 15 min (excitation 485 nm, emission 528 nm). Droplets were incubated for between 30 and 40 h. An Olympus DP71 microscope was used to examine the cell growth.

#### 2.6.2 Droplet sorting

Following off-chip incubation, droplets were poured into a capped syringe (Global, 1 mL) and any remaining volume of the syringe was filled with fluorinated oil with 2% surfactant before inserting the syringe plunger. We reinjected the droplets into a droplet detection/sorting device with height 50 μm and main channel width 55 μm using a syringe pump (KD Scientific). The syringe was connected to the sorting device via a 23-gauge needle and 0.022” ID PTFE tubing. Droplets of 55 μm diameter were reinjected into the sorting device at 1.5-2.5 μL/min, corresponding to ~150-300 droplets/sec. Sorting was performed in a manner similar to that of (Chung et al. 2017), with the major difference being that droplets were generated using a droplet generation device (Fig. S9A), incubated off-chip to allow cell growth, and then reinjected into a sorting device (Fig. S9B). After sorting, the cell identity was determined either by the fluorescence of the cells within the droplet, or by the colony color after chemically destabilizing the droplets and plating on MacConkey agar with galactose. Both the sorting method and strain identification are described in more detail in the Supplementary Information.

## 3. Calculation: Model-based prediction of amplification of production improvement via metabolic cross-feeding circuits

Kerner et al. (2012) presented an ODE model of a cross-feeding co-culture in its exponential growth phase, assuming constant secretion and uptake parameters and Monod kinetics for growth on a limiting nutrient (i.e., the amino acid for which the strain is auxotrophic). In this model, the co-culture reaches a pseudo-steady-state in which the two strains have the same growth rate (*μ*, unit: 1/hr), which depends on each auxotroph’s secretion rate (*α*_Sec_, *α*_Sens_; unit: mmol/gDM-hr) of its shared metabolite and the per cell growth requirements for the cross-fed metabolites (*β*_Sec_, *β*_Sens_; unit: mmol/gDM). Additionally, a steady population composition ratio (r) between the number of each cell type (*N_Sec_*, *N_Sens_*) is reached. Mathematically, these properties are given by:

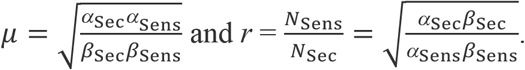

Thus, if the secretion rate of the secretor (*α*_Sec_) increases and all else remains unchanged, we expect the co-culture growth rate to increase and the percentage of the final population that is the sensor to increase. If we have a base-level secretor strain with secretion rate *α_Base_* and an improved secretor with secretion rate *α*_Base_ (1+*x*) and we grow each secretor with the sensor strain, the exponential growth of the co-culture quickly amplifies even moderate differences in production level (Fig. 2). After an amount of time corresponding to *n* doublings of the base strain co-culture, the improved secretor’s co-culture will have 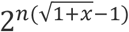 times as many cells as the base strain’s co-culture. For comparison, traditional methods of growing an auxotrophic biosensor on the production strains’ supernatants would result in 1 + *x* times as many cells on the higher producer’s supernatant as on the base strain’s, with the assumption that the higher producer accumulates 1 + *x* higher product concentration and the concentrations are within the biosensor’s linear range. For example, an improved production strain with a 50% increase in production rate can result in 375% more cells in the co-culture containing the improved secretor than in that with the base strain, corresponding to an amplification of 7.5 (Fig. 2, point A). In contrast, a monoculture of the auxotrophic biosensor on the supernatant from these strains would produce merely 50% more cells for the high-producer than for the base, exhibiting no amplification.

**Figure 2:**
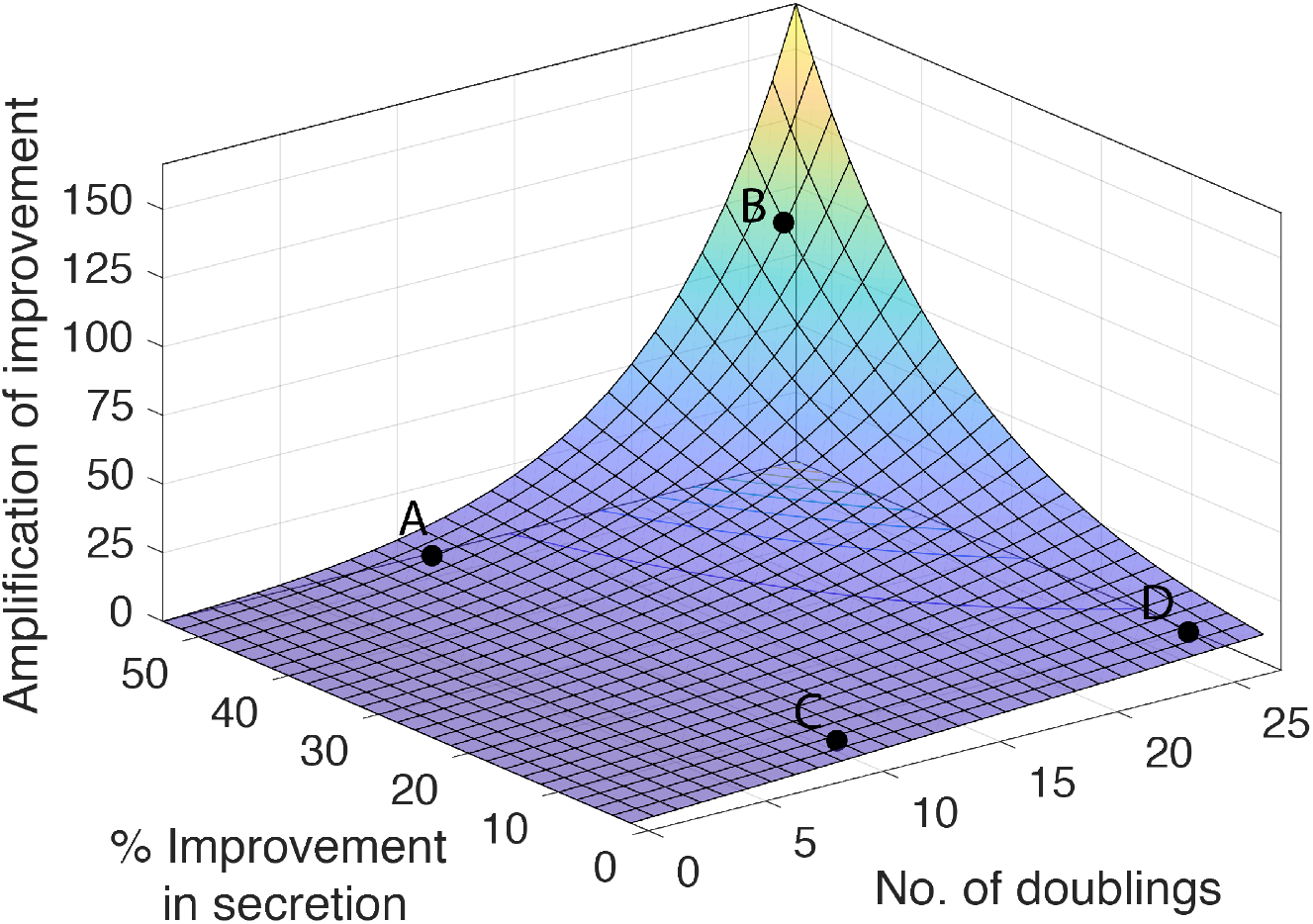
Amplification of secretion improvement by co-culture growth. A production strain with 50% improvement compared to a base strain will lead to a co-culture with 4.75 times as many cells as that of the base strain (375% more) after a time corresponding to 10 doublings of the base co-culture. This is a 7.5-fold amplification of the percentage increase (point A). After 25 doublings of the base strain co-culture, this amplification will rise to 96-fold (point B). For a smaller improvement of 5%, there will be 1.19 times as many cells in the improved secretion co-culture after 10 base doublings, a 3.7-fold amplification (point C). After 25 doublings of the base co-culture, this will increase to an 11-fold amplification (point D).

At some high enough *α*_Sec_*N*_Sec_ such that the target molecule is no longer limiting for the sensor strain, this model breaks down. There is then one-directional feeding where the sensor can grow without further growth of the secretor strain. In this case, the maximum growth rate may not be increased for a higher-producing strain, but the time to reach this critical value of *α*_Sec_*N*_Sec_ will vary for secretor strains with different *α*_Sec_ production rates. We, therefore, expect that, over some range of secretion levels, improvements in secretor strain production will lead to detectable changes in co-culture growth and composition.

## 4. Results and discussion

### 4.1 Development of a cross-feeding circuit for screening 2-ketoisovalerate production

We will demonstrate the SnoCAP screening framework using 2-ketoisovalerate (2-KIV) as a target molecule. 2-KIV is a precursor of the branched-chain amino acids valine and leucine. By overexpression of an alpha-ketoisovalerate decarboxylase (Kdc) and an alcohol dehydrogenase (Adh) in *E. coli*, 2-KIV can be converted into the drop-in biofuel isobutanol. Additional overexpression of three enzymes that catalyze the conversion of pyruvate to 2-KIV leads to substantially improved isobutanol production (Atsumi, Hanai, and Liao 2008) (Fig. 3A,B). We chose a Δ*ilvD* auxotroph as the sensor strain. IlvD, dihydroxy-acid dehydratase, catalyzes the conversion of 2,3-dihydroxy-isovalerate into 2-KIV. IlvD is also part of the isoleucine biosynthesis pathway, catalyzing the conversion of 2,3-dihydroxy-3-methylvalerate into 2-keto-3-methylvalerate. We therefore supplemented the co-cultures with an excess of isoleucine in order to eliminate effects from variation in isoleucine cross-feeding levels. When grown with excess isoleucine and varying levels of 2-KIV, the Δ*ilvD* auxotroph’s growth rate and maximum cell density increase in response to increasing 2-KIV over a certain range (Fig. S1A,C).

For a given *E. coli* auxotroph, a variety of cross-feeding partner auxotroph options are generally available, exhibiting a range of co-culture growth rates (Wintermute and Silver 2010; Mee et al. 2014). We tested a panel of potential partner auxotrophic strains for their ability to cross-feed with K12 Δ*ilvD* in a minimal medium supplemented with isoleucine. Of the partner auxotrophs tested (Δ*hisD*, Δ*leuB*, Δ*lysA*, Δ*pheA*, Δ*ppC*, Δ*trpB*, and Δ*tyrA*, each in strain BW25113), Δ*lysA* (a lysine auxotroph) and Δ*pheA* (a phenylalanine auxotroph) showed consistent growth (Fig. S2). Both showed considerable lag phases (~2 and 5 days, respectively) and lower maximum optical densities than monocultures, leaving room for improved secretion to boost co-culture growth. We chose to proceed with lysine as the secondary cross-fed molecule. Selecting a different auxotroph, including ones that have no growth with the base production-level strain, may be a useful strategy to adjust the dynamic range of the screening.

We tested the growth properties in 96-well microplates of sensor strain K12 Δ*ilvD* in co-culture with secretor strains of several different secretion levels. Genotypes of these strains are listed in Table S1. JCL16 Δ*lysA* is the base strain, with a low production level. JCL260 Δ*lysA alsS*, which has six gene deletions to direct flux through the isobutanol pathway and a single copy of *alsS*, under an IPTG-inducible promotor, integrated into the genome, represents an intermediate-production-level strain. JCL260 Δ*lysA* pSA69, which has the same gene deletions and carries a plasmid for overexpression of *alsS* and *ilvCD* under an IPTG-inducible promotor, represents a high-production-level strain. In monoculture fermentations, these strains have differing production levels of 2-KIV (Fig. 3B). When transformed with pSA65, which carries *kivD/adhA*, the strains have differing levels of isobutanol production and glucose consumption, and similar growth profiles (Fig. 3C). We inoculated these strains (lacking pSA65) in co-culture with the sensor strain at a 1:1 initial ratio in minimal medium with isoleucine. Monoculture inoculation of any of these strains or the sensor strain in this medium produces no detectable growth. In the co-cultures, growth order (Fig. 3D, E) and growth rate (Fig. 3F) increases with increasing strain production level. Addition of IPTG results in a higher growth rate for co-cultures with each of the strains carrying an IPTG-inducible operon (either *alsS* only or *alsS-ilvCD*), and no difference for base strain JCL16 (Fig. 3D-F). This indicates that expression of the genes leading to 2-KIV translates into increased growth in the co-culture setting. We determined the composition of the co-cultures when in early stationary phase by differential plating (see Supplementary Methods). As predicted, the sensor-to-secretor strain ratio rises with increasing production level of the secretor strain (Fig. 3G).

**Figure 3:**
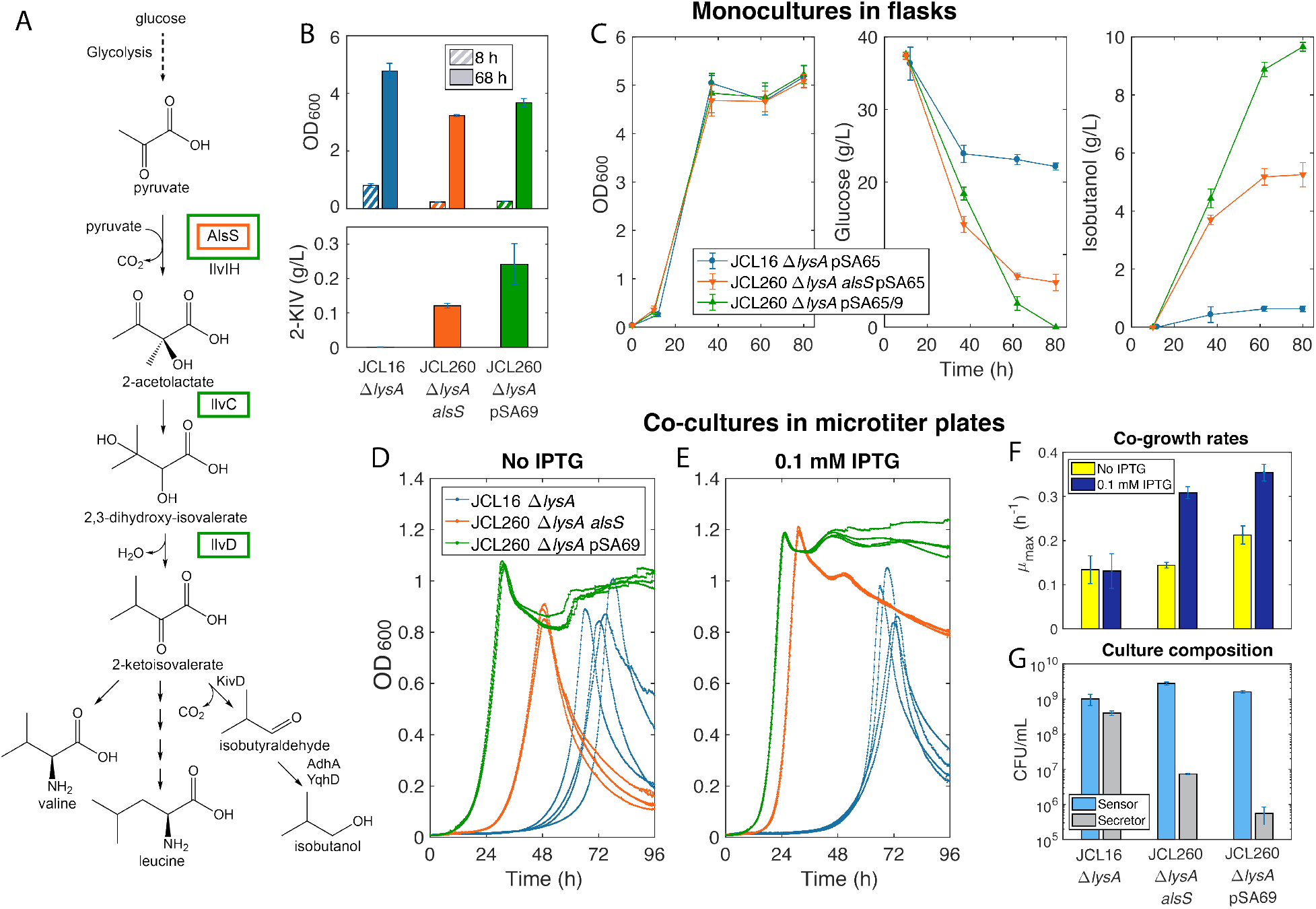
Co-culture growth characteristics correlate with monoculture production performance. (A) 2-KIV and isobutanol production pathway. Genes that are overexpressed in the production strains are boxed in the color corresponding to the strain in the legend in (C). (B) Monoculture 2-KIV production characteristics of three different secretor strains on 20 g/L glucose. IPTG was added after 8 h and 2-KIV was measured 60 h after induction. Error bars represent the standard deviation of three biological replicates. (C) Monoculture isobutanol production characteristics of these three strains when carrying pSA65 P_L_lacO_1_::*kivd–adhA*). Isobutanol, glucose and cell growth profiles are shown. Error bars represent the standard deviation of four biological replicates. Growth profiles of co-cultures of the same three secretor strains (not carrying pSA65) with sensor strain K12 Δ*ilvD::kan* in a microplate with (D) and without (E) IPTG-induced expression of the *alsS/ilvCD* genes. Three replicate wells are shown for each strain pair, plotted in the same color. (F) Co-culture growth rates. Error bars represent the standard deviations between at least 10 biological replicates, combined from two separate experiments. (G) Population composition of microplate cultures in early stationary phase. Error bars represent standard deviations from two biological replicates with two technical replicates each.

To evaluate the suitability of the cross-feeding co-culture growth assay for high-throughput screening, we calculated the Z-factor, a parameter that reports on a combination of the signal dynamic range and assay precision (Zhang, Chung, and Oldenburg 1999). Values between 0.5 and 1.0 are considered indicative of an excellent assay. We considered co-cultures containing secretor JCL16 Δ*lysA* as the negative control and JCL260 Δ*lysA* pSA69 as the positive control and found that the Z-factor was in the “excellent” range from hours 11 to 62 of the assay period (Fig. S3).

### 4.2 Implementation of co-culture screening in an agar plate format

We next implemented SnoCAP as a colony-screening assay. By spreading ~10^7^ CFU of the sensor strain (enough to form a lawn if the cells were able to grow in monoculture) and ~100 CFU of secretor strain on agar medium in 10 cm diameter Petri dishes, we obtained mixed colonies, each originating from a single secretor cell. We verified that these colonies were indeed mixtures of secretor and sensor by streaking them on LB plates with IPTG/X-gal and observing both blue (sensor) and white (secretor) colonies. We also observed no growth on monoculture plates containing only one strain, demonstrating that colonies only form when both sensor and secretor are present. We compared mixed colony formation between the base strain (JCL16 Δ*lysA*) and the high-producing strain (JCL260 Δ*lysA* pSA69). JCL16 colonies appeared later and ultimately formed flatter, more translucent mixed colonies that are easily distinguishable from the taller, opaque JCL260 Δ*lysA* pSA69 mixed colonies, even once both have grown to a substantial footprint area (Fig. 4A). Higher-production-level secretors also produced colonies with increased sensor-to-secretor ratio (Fig. 4B).

**Figure 4:**
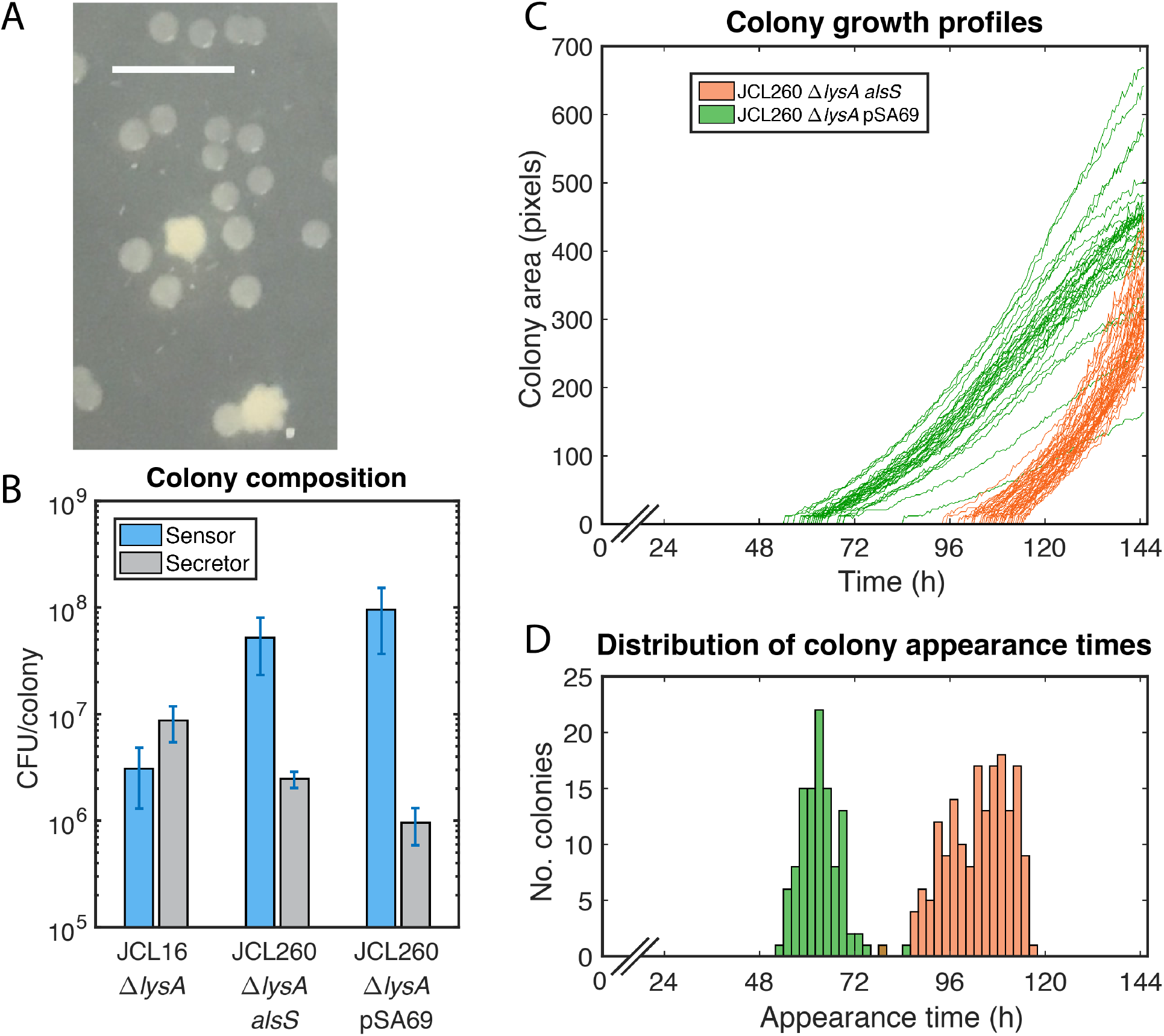
Implementation as a colony screening assay on agar plates. (A) Mixture of JCL16 Δ*lysA* and JCL260 Δ*lysA* pSA69 mixed colonies after 6 days at 37 °C. More opaque white colonies contain JCL260 Δ*lysA* pSA69, while more translucent colonies are JCL16 Δ*lysA*. Scale bar represents 0.5 cm. (B) Composition of colonies after 7 days incubation at 35 °C. Error bars represent standard deviations of 4 colony replicates, with 4 technical replicates each. (C) Colony area over time for the plates. Growth curves are shown from 55 colonies on a JCL260 Δ*lysA alsS* plate and 48 colonies on a JCL260 Δ*lysA* pSA69 plate. (D) Histogram of colony appearance times, combined from three plates of each secretor strain (126 total colonies from JCL260 Δ*lysA* pSA69 plates and 110 colonies from JCL260 Δ*lysA alsS* plates).

We next compared the growth of the intermediate level strain (JCL260 Δ*lysA alsS*) and the high production strain (JCL260 Δ*lysA* pSA69). Here, both strains eventually formed large opaque colonies, but JCL260 Δ*lysA alsS* colonies appeared later. We implemented the ScanLag technique for automated imaging and colony growth profile analysis developed by Levin-Reisman et al. (Levin-Reisman et al. 2010; Levin-Reisman, Fridman, and Balaban 2014), to observe colony lag time and growth dynamics. This inexpensive method uses photo scanners to image the Petri dishes periodically during growth and a MATLAB-based application that aligns the images and returns colony growth phenotype information. The analysis revealed that JCL260 Δ*lysA alsS* mixed colonies appear later and then grow to eventually reach similar colony size as JCL260 Δ*lysA* pSA69 mixed colonies (Fig. 4C, D). This quantification of colony size and dynamics helps to reduce subjectivity in assessing positive hits from the screen.

### 4.3 Increasing the dynamic range by utilization of an inhibitory analog of the target molecule

One common issue in previous biosensor-based screening is the limited dynamic range of production levels over which the biosensor responds. We hypothesized that the addition of an inhibitory analog of the target molecule could expand the dynamic range of screening (i.e., the concentrations of 2-KIV over which the co-growth changes in response to increases in 2-KIV) and increase differences of co-culture growth in the SnoCAP framework. Analog selection, in which bacteria are grown on an inhibitory analog of a metabolite in order to select for overproduction of that metabolite, is a useful strategy for strain development and has been used to identify mutations that overexpress pathway genes or decrease feedback inhibition from target molecules (Karlström 1965; Valle and Berry 1999). Metabolite analogs exist for various types of compounds, including amino acids, vitamins, organic acids, and sugars. A computational framework has recently been developed to identify candidate metabolite analogs for use in strain improvement (Cardoso et al. 2018).

Norvaline, a toxic analog of valine and leucine, has been used as a selection agent for increased flux through the valine (Karlström 1965) and isobutanol (Smith and Liao 2011) pathways. Norvaline produces a growth defect that can be partially recovered by the addition of either leucine or valine alone or fully recovered by addition of both leucine and valine (Fig. S4). We first verified that the sensor strain’s growth remains responsive to increasing 2-KIV in the presence of norvaline (Fig. S1B, D). We then added norvaline to the co-cultures and found that its addition magnifies differences between the strains at the higher end of the production spectrum (Fig. 5A). For low levels of norvaline, the co-cultures with JCL260 Δ*lysA* pSA69 as the secretor strain are not affected by the norvaline, while those with JCL260 Δ*lysA alsS* exhibit an increasing lag phase time with increasing norvaline concentration. For higher levels of norvaline, such as 1.0 g/L, co-cultures with JCL260 Δ*lysA* pSA69 show lengthening lag phase as well. Thus, norvaline can be used to expand the dynamic range and increase the production threshold below which secretor strains cannot support co-culture growth.

**Figure 5:**
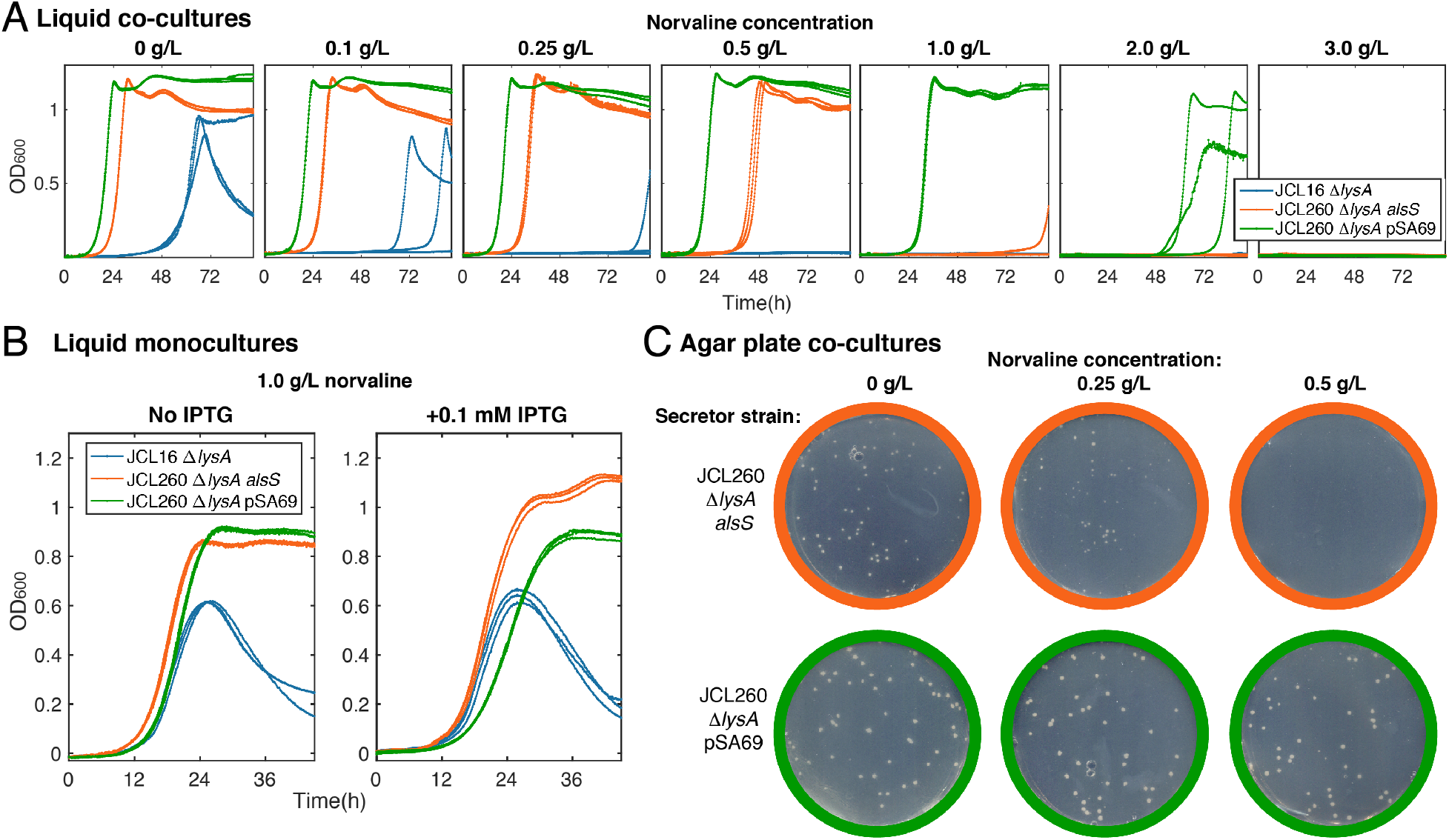
Tuning dynamic range of the 2-KIV screening by addition of norvaline. (A) Liquid co-cultures with various levels of norvaline. (B) Liquid monocultures on 1.0 g/L norvaline, demonstrating that norvaline is not an effective selection tool for these strains in monoculture. (C) Agar plate co-cultures containing various concentrations of norvaline, after 168 h incubation at 35 °C. At 0.5 g/L norvaline, JCL260 Δ*lysA alsS* plate has no visible colonies.

It is also interesting to note that the co-culture growth is not strongly affected by factors that may govern monoculture growth, such as over-expression of synthetic gene constructs. For instance, although pSA69 increases 2-KIV production, we observed that induction of its expression with IPTG results in decreased monoculture growth in norvaline-supplemented medium (Fig. 5B). Yet in norvaline-supplemented co-cultures, JCL260 Δ*lysA* pSA69 produces the fastest co-growth. This supports the prediction of our model that co-culture growth characteristics are determined primarily by the production levels of the cross-fed metabolites.

In the agar plate assay, norvaline addition was also effective in widening the difference in growth between co-cultures containing the intermediate- and high-producing secretor strains (Fig. 5C). At 0.5 g/L norvaline, only the highest producing strain (JCL260 Δ*lysA* pSA69) formed opaque colonies, while the intermediate-producer (JCL260 Δ*lysA alsS*) developed smaller, translucent colonies that were not visible in the scanner images.

To evaluate the ability of the agar plate screening assay to identify rare higher-producing strains, we tested model libraries consisting of the intermediate-producing strain spiked with a small percentage (0.6% or 0.1% of the population) of the high-producing strain. On each plate we observed the development of many small translucent colonies as well as a smaller number of large opaque colonies. We isolated the secretor from the large colonies and investigated the strain identity. For the 0.6% library, all of the 25 large colonies except one were identified as containing the high-secreting strain. The remaining one was found to be a colony containing both the intermediate- and high-producing secretor strains. For the 0.1% library, three large colonies emerged and were all true positives (for more details on the model libraries, see Fig. S5 and Supplementary Note 1). We concluded that, when an appropriate level of norvaline is utilized for the range of strain improvement that is targeted, the screening can easily identify rare higher production level strains at frequencies as low as 0.1% (or lower if a larger number of plates are employed). Transformation with pSA65 and production testing of secretor cells isolated from the colonies confirmed that the strain is not adversely affected by the screening conditions (Fig. S5C).

### 4.4 Development of a cross-feeding co-culture system for screening L-tryptophan production

As a second test case of the SnoCAP screening framework, we examined tryptophan-producing strains. For this implementation, we chose BW25113 Δ*trpB*, which lacks the catalytic subunit of tryptophan synthase, as the sensor strain. We selected histidine as the secondary crossfed molecule (Fig. 6A), based on work showing that growth of tryptophan/histidine cross-feeding co-cultures is affected by overproduction of tryptophan (Pande et al. 2015). For overproduction strains, we deleted the *trpR* and *tnaA* genes, both individually and sequentially in the same strain. Deletion of these genes, which encode a repressor of the *trp* operon and a tryptophanase, respectively, are typical early steps in the engineering of tryptophan production strains (e.g., in the work of (Carr et al. 2012; Zhao et al. 2011)). In liquid co-cultures in microplates, all three modified strains had increased co-culture growth compared to the base strain. However, the growth took several days to be observable. Since these deletions are only first steps in tryptophan strain engineering, it may be useful that these strains’ co-growth is slow, since it leaves substantial room for improvement with higher production-level strains. Nevertheless, we were also interested in determining whether we could decrease assay time while still maintaining the detectable differences in growth phenotype between the higher production-level strains and base strain. We added low levels of histidine and tryptophan to jump-start the co-cultures and found that addition of histidine produced the desired effect (Fig. 6A). Addition of tryptophan does not have a noticeable effect, indicating that histidine levels limit the initial growth, with growth not commencing until the initial secretor strains have secreted enough histidine. This is not unexpected since the secretor strains overproduce tryptophan, whereas the sensor strain does not overproduce histidine. Engineering the production level of the secondary cross-fed molecule by the sensor strain may be an effective approach for tuning the assay timing and dynamic range.

We compared the Δ*trpR* strain and the base strain in the agar plate-screening assay. Similar to the microplate, colonies were slow to develop on plates that were not supplemented with any amino acids, but after two weeks we observed colonies on the Δ*trpR* secretor co-culture plates and none on the base secretor co-culture plates, and none on any monoculture plates. Addition of small initial amounts of histidine produced a significant decrease in time required for colony visibility while maintaining clear differences in the growth characteristics between the two secretor strains (Fig. 6B, C). In this case, both the base and improved secretor strains formed colonies, but the base strain colonies were flat, translucent, and easily distinguishable from the taller, opaque *DtrpR* secretor colonies.

**Figure 6:**
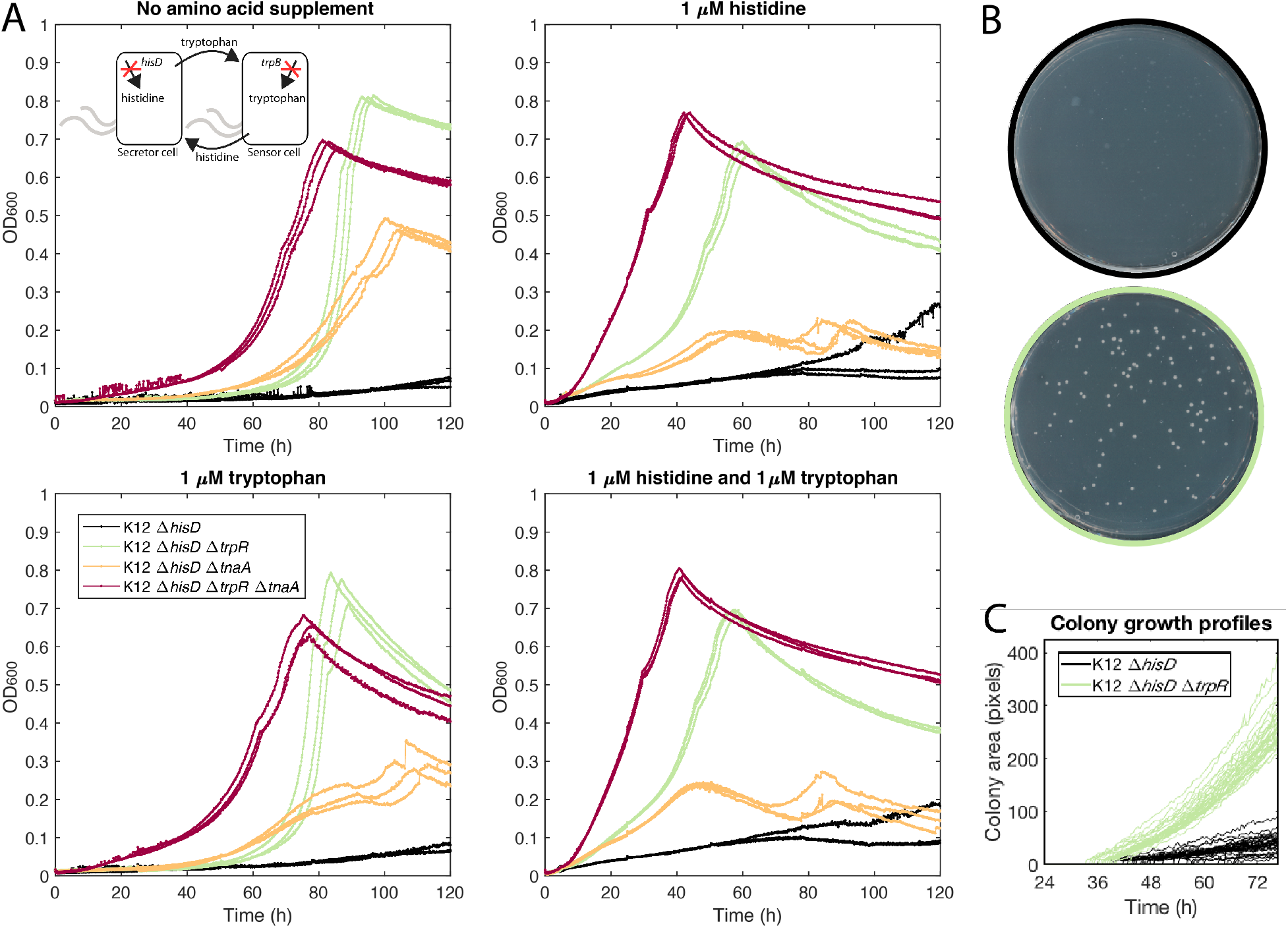
Implementation of the screening method for tryptophan production. (A) Co-culture growth in 96-well microplates with different initial amino acid supplementation. Three biological replicates are shown for each strain, plotted in the same color. The inset shows the scheme of tryptophan/histidine cross-feeding. Images after 72 h incubation (B) and colony growth profiles (C) of agar plate co-cultures comparing secretor strains K12 Δ*hisD* and K12 Δ*hisD* Δ*trpR*, with 5 μM initial histidine.

### 4.5 Application of an intermediate-sensor assisted push-pull strategy to screen for isobutanol production

Intermediate-sensor assisted push-pull strategy has been proposed as a general method for screening for production of molecules that lack a direct sensor but for which a sensor is available for an intermediate, and has been successfully demonstrated for deoxyviolacein with a tryptophan biosensor (Fang et al. 2016). This approach involves using the intermediate sensor to screen for increased production of the intermediate and then improving the conversion of intermediate to final product by screening for decreased readout from the biosensor. We investigated whether SnoCAP can be employed in such a manner, i.e., whether increased activity of a pathway converting the cross-fed molecule to a target molecule will decrease the co-culture growth.

Using the 2-KIV system, we compared co-culture growth with secretor strain JCL260 Δ*lysA* pSA69, a strain in which the pyruvate to 2-KIV part of the pathway performs well, to that of secretor strain JCL260 Δ*lysA* pSA65/9, which additionally carries the 2-KIV to isobutanol part of the pathway. We observed a significant decrease in co-culture growth rate and increase in lag time for the pSA65 carrying strain (Fig. 7A). We also examined intermediate levels of KivD/AdhA expression in strains that contain copies of these genes integrated into the genome. These strains were generated using chemically inducible chromosomal evolution (CIChE), which enables the copy number of a construct of interest to be adjusted by changing the concentration of a lethal chemical (e.g., chloramphenicol, to which the resistance can be rendered by the integrated construct in a dose-dependent manner) in the medium (Tyo, Ajikumar, and Stephanopoulos 2009). Using this method, we obtained strains JCL260 Δ*lysA* cm 20 and JCL260 Δ*lysA* cm 80. These strains have differing levels of isobutanol production (Fig. 7B). We saw that with the lower levels of Kdc/Adh expression there is still a decrease in co-culture growth but to a lower degree than with the high expression from pSA65 (Fig. 7C). We observed similar trends when testing these strains in the agar plate assay, with the mixed colonies appearing later for secretor strains with higher Kdc/Adh expression levels (Fig. 7D, E). We further verified that the decreased co-culture growth with the pSA65-carrying strain was indeed due to 2-KIV being channeled away from cross-feeding, instead of other factors such as metabolic burden from overexpression of the plasmid genes or the toxicity of the isobutanol product, by employing conditions that force cross-feeding independent of 2-KIV (see Fig. S6 and Supplementary Note 2). These results indicate that the push-pull strategy can be effectively applied to this system with a dynamic range that includes expression levels that are relevant for strain engineering.

### 4.6 Screening of a chemically-mutagenized strain library for improved plasmid-free isobutanol production

We next applied the SnoCAP screening method to strain development for higher 2-KIV production based on genomic modifications (rather than by pathway overexpression from the pSA69 plasmid). We introduced random mutations into the genome of JCL260 Δ*lysA alsS* by NTG-mutagenesis. We plated the library on 24.5 cm square Petri dishes containing various concentrations of norvaline, along with an excess of sensor strain. ~10^4^ LB-CFU (cells that would form colonies in monoculture on an LB plate) of secretor strain library were spread on each plate. After seven days of incubation at 37 °C, we identified large opaque colonies among the large number of small translucent colonies. We isolation streaked large colonies from the 1.0 g/L norvaline condition on LB plates with tetracycline to separate the secretor strains and then rescreened them in the microplate co-culture assay and selected 7 isolates that showed improved co-culture growth. We transformed these isolates with pSA65 and tested their isobutanol production. Due to reduced growth rates observed in some of the library isolates, we tested their production levels in M9IPG supplemented with 5 g/L yeast extract. Of the 7 isolates tested, one, which we call strain B1, showed significantly improved production compared to the base strain. After 73 h fermentation, B1 pSA65 produces 9.4 ± 0.4 g/L (SD, n = 3) isobutanol, representing 60% of the theoretical yield, compared to 1.8 ± 0.1 g/L and 16% theoretical yield by the parental strain JCL260 Δ*lysA alsS* (Fig. S7). It should be noted that JCL260 Δ*lysA alsS* performs less well under yeast extract supplemented conditions compared to minimal medium conditions, likely due to lower expression of the *ilvCD* genes when it is not necessary to produce all of its own amino acids. Nevertheless, B1 isolate performs superiorly even to JCL260 Δ*lysA* alsS’s production level in minimal medium (i.e., the production level presented in Fig. 5B). B1 performs nearly as well as JCL260 Δ*lysA* pSA69, but through a different mechanism than plasmid overexpression of the *ilvCD* genes. To investigate the mutations leading to this improved production, we sequenced the genome of B1 and parental strain with >100X coverage and identified 73 SNPs, 39 of which lead to amino acid substitutions or stop codon introductions, in B1 (Dataset S1). One intriguing mutation occurs in the *aceK* gene, which encodes the isocitrate dehydrogenase kinase/phosphatase. This bifunctional enzyme controls the branch between the TCA cycle and glyoxylate cycle by modification of isocitrate dehydrogenase. B1’s mutation consists of a proline to serine substitution in residue 510, which is part of the substrate recognition loop (Zheng and Jia 2010). We reintroduced this mutation into a *mutS-* version of the parental strain by singlestranded oligo recombination, calling this strain JCL260 Δ*lysA alsS aceK-mut*. When tested in liquid co-culture with the sensor strain, JCL260 Δ*lysA alsS aceK-mut* shows improved co-growth compared to the parental strain (Fig. S8A). After transformation with pSA65, we compared the production and found that this mutation does, indeed, lead to a modest increase in isobutanol production (Fig. S8B).

**Figure 7:**
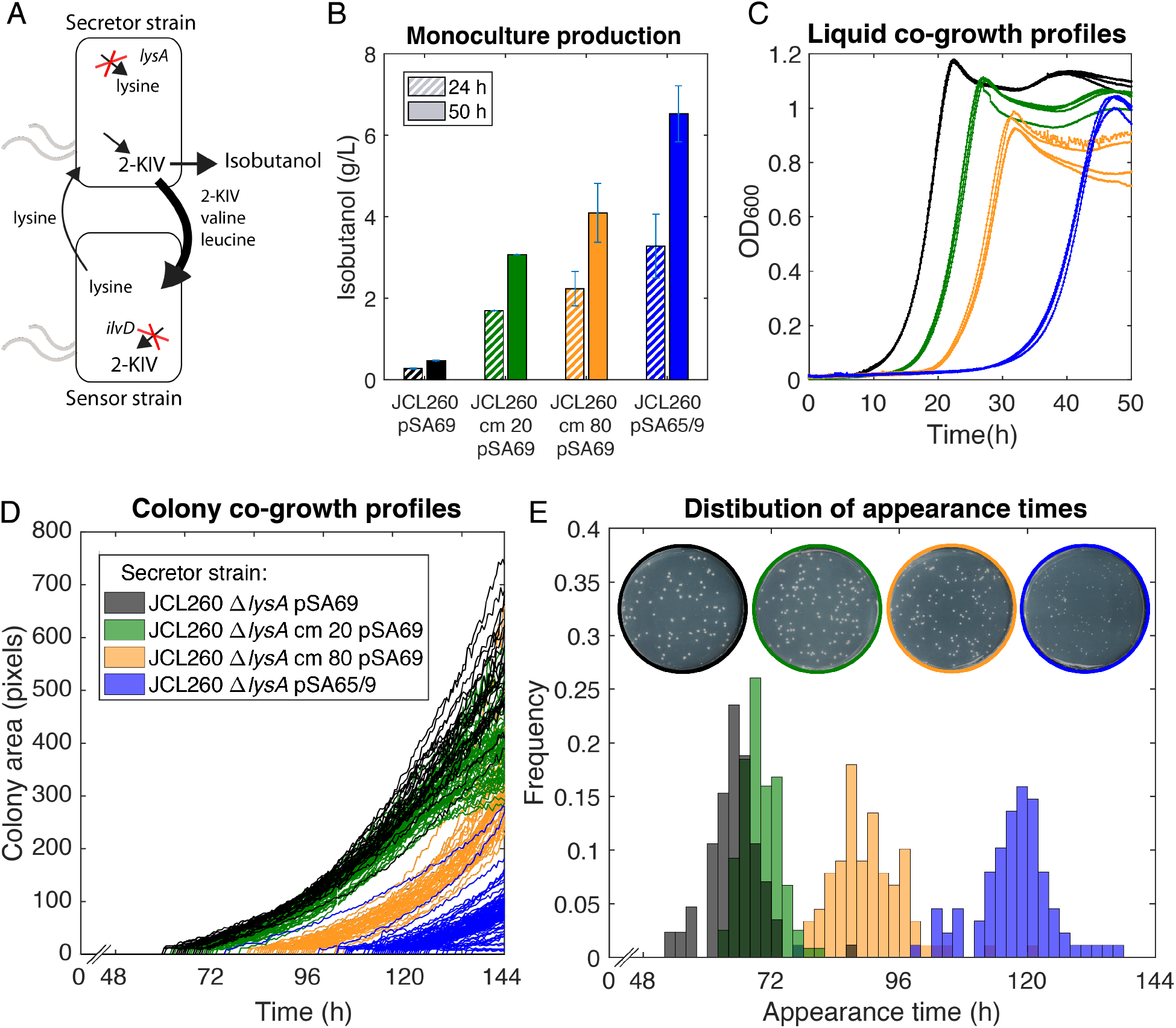
Screening for conversion of 2-KIV into isobutanol. (A) Schematic of sensing intermediate. (B) Production levels of the four strains with differing levels of *adhA/kivD* expression in M9 minimal medium, 36 g/L glucose. (C) Liquid co-culture growth profiles of secretor strains carrying varying levels of *adhA/kivD* expression, colony growth profiles (D) and colony appearance time histogram (E) for agar plate assay comparing the four secretor strains. Plot in (F) shows profiles for the unmerged colonies from one plate of each secretor strain (29, 50, 34, 84 for JCL260 Δ*lysA* pSA69, JCL260 Δ*lysA* cm 20 pSA69, JCL260 Δ*lysA* cm 80 pSA69, JCL260 Δ*lysA* pSA65/9, respectively) (the JCL260 Δ*lysA* pSA65/9 plates have more unmerged colonies because colony size is smaller). The histogram in (E) shows appearance times for the unmerged colonies from a combination of two plates from each strain except for JCL260 Δ*lysA* pSA65/69 for which colonies from one plate are included since smaller colony size resulted in reduced merging and more analyzable colonies per plate. In total, this amounts to 85 from JCL260 Δ*lysA* pSA69, 119 from JCL260 Δ*lysA* cm 20, 89 from JCL260 Δ*lysA* cm 80, and 84 from JCL260 Δ*lysA* pSA65/9. Plate images above the histogram in (E) were taken after 144 h incubation. Legend in (D) also applies to (C) and (E).

### 4.7 Implementation of cross-feeding co-culture screening via high-throughput microdroplet cocultivation and sorting

To further increase the throughput of the SnoCAP screening framework, we next investigated compartmentalization by encapsulation in microfluidic water-in-oil droplets. These monodisperse droplets, with volumes in the picoliter to nanoliter range, provide miniaturized culture volumes that can be analyzed in a variety of ways, including by high-throughput automated sorting to isolate droplets containing the highest fluorescence signal. Cells are distributed according to a Poisson distribution, and cell density can be manipulated to ensure that initially i) all droplets contain several sensor cells, and ii) the majority of droplets contain either zero or one secretor cell.

Droplet sorting, either via microfluidic sorting devices or commercial flow cytometry, is emerging as a valuable tool for screening strain libraries at high-throughputs (Baret et al. 2009; Wang et al. 2014; Huang et al. 2015; Siedler et al. 2017; Abatemarco et al. 2017; Wagner et al. 2018; Li et al. 2018). With current technology, the highest throughput of sorting is achieved using fluorescence signal. To couple the co-growth output to fluorescence, we expressed fluorescent proteins in the strains. We first labeled the high secretor, JCL260 Δ*lysA* pSA69, with YFP, the sensor strain with mCherry, and observed co-culture growth in droplets (Fig. 8A). For sorting experiments, we employed a different version of the sensor strain carrying a plasmid encoding constitutively expressed mNeonGreen and screened by fluorescence-activated droplet sorting (FADS) for droplets with the highest fluorescence, corresponding to the largest number of sensor cells.

We tested the ability of FADS to distinguish between two strains of differing production levels and to isolate the high secretor, JCL260 Δ*lysA* pSA69, when spiked at low percentages into a population of lower secretor cells. The fluorescence signal profiles of monosecretor-controls indicated that the higher-producing strains result in droplets with more fluorescence and that thresholds can be established to separate the different production level secretor strains (Fig. 8B). We identified conditions (i.e., an appropriate norvaline concentration) that enabled the desired level of separation between the two strains (Fig. 8B), encapsulated model libraries consisting of mixtures of the two secretor strains (device shown in Fig. S9A), incubated to allow co-growth with the sensor strain within the droplets, and then sorted to isolate the most fluorescent droplets. We mixed JCL260 Δ*lysA* pSA69 and JCL260 Δ*lysA alsS* Δ*galK* (*galK* deletion enabled color-based differentiation of the strains when plated on MacConkey agar with galactose, Fig. S11B) at a ratio of 1:100 and encapsulated them with sensor strain K12 *ilvD::kan* pSAS31 on 1.75 g/L norvaline. We used cell densities such that the secretor cell loading was ~0.1/droplet and sensor cell loading of ~5/droplet. Cells are encapsulated according to the Poisson distribution and these loadings lead to most droplets containing either no secretor cell or a single secretor cell. Although ~90% contain no secretor, using a higher secretor loading will lead to more co-encapsulation of two secretor cells in the same droplet and hence more false positives. The sensor cell loading ensures that almost all droplets contain at least one sensor cell while leaving room for growth.

**Figure 8:**
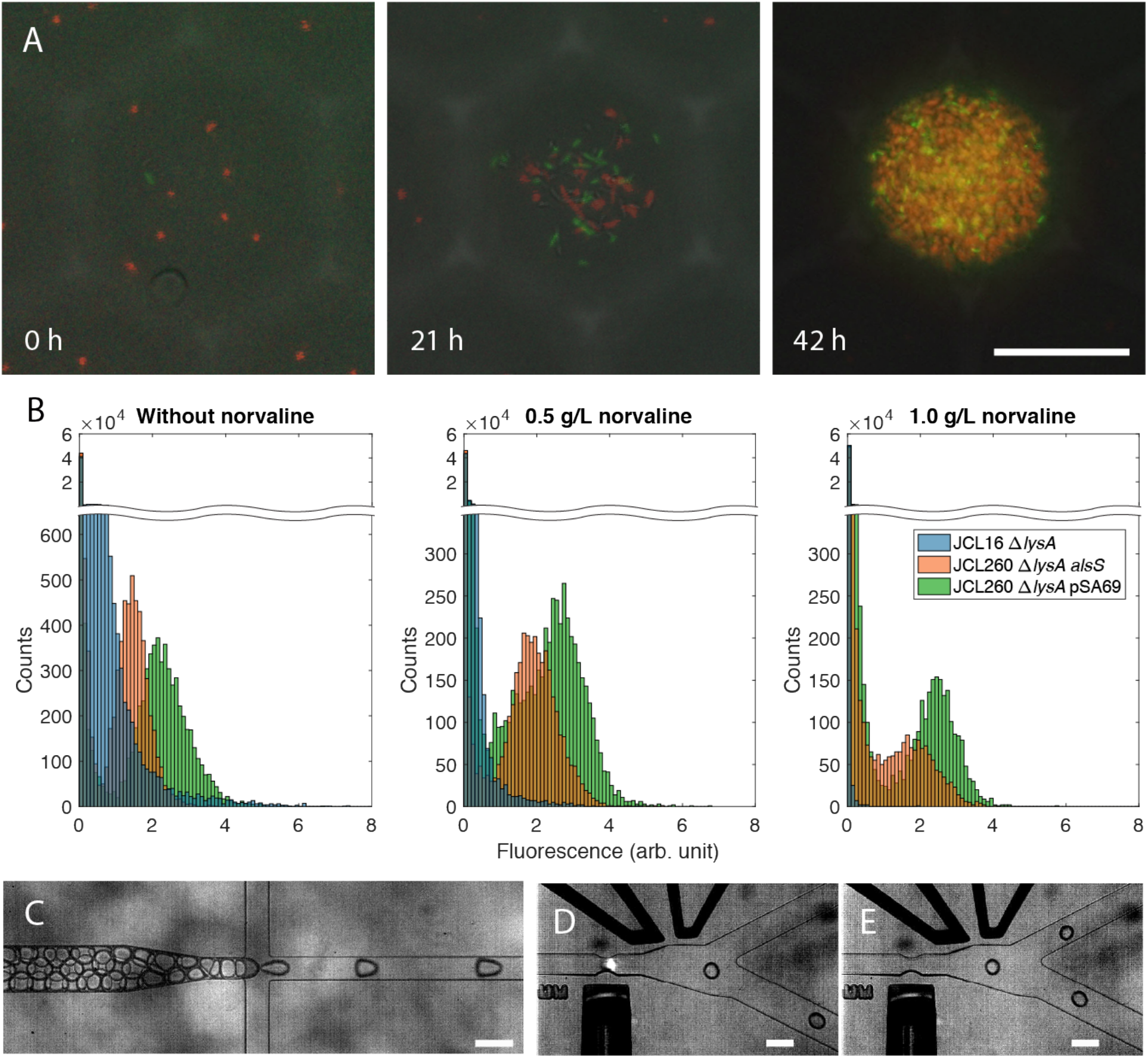
Screening for 2-KIV production implemented in microfluidic droplets. (A) Representative droplet images during co-growth of sensor strain expressing mCherry and JCL260 Δ*lysA* pSA69 expressing YFP on 1.0 g/L norvaline. Each image is an overlay of fluorescence and brightfìeld images, but cell locations do not align exactly due to cell movement between images. (B) Histograms comparing fluorescence signal from droplets containing co-cultures with different secretor strains (at an initial cell loading of 0.2 secretor and 5 sensor/droplet) on several concentrations of norvaline after a 30 h incubation. ~50,000 droplets under each condition were analyzed. (B) Droplet reinjection and spacing. (C, D) Fluorescence-activated droplet sorting (FADS) to retrieve droplets with high levels of sensor strain growth. Scale bars: (A) 50 μm, (C-E) 100 μm.

Following incubation to allow co-culture growth, the droplets were reinjected into a sorting device (Fig. 8C, Fig. S9B, Movie S1) and sorted based on fluorescence from excitation with a 450 nm laser (Fig. 8D, E, Movie S2). We selected sorting gate values based on comparisons of the signal profiles from mono-secretor control sets of droplets, and verified their effectiveness by observing that these values enabled bright droplets to be collected. For application to a library when a high-producing strain is not already available, the threshold can be selected to sort the top percentage of droplets at a desired stringency. After sorting, desired droplets were collected into a PDMS device with an elevated chamber to retain droplets while allowing oil to flow out (Fig. S10). In a 25 min sorting period, we isolated 10 droplets, each of which showed significant sensor cell growth. Plating these droplets on a MacConkey agar plate with tetracycline (to prevent growth of the sensor strain) produced only purple colonies (>100- fold enrichment of the high-secretor) (Fig. S11D), whereas plating the unsorted droplets produced a mixture with 16% purple colonies (16-fold enrichment) (Fig. S11C). We assessed the productivity of purple colonies that resulted from sorted droplets after transformation with pSA65 and found that they maintained their productivity through the screening process (Fig. S12).

We noticed lower than expected cell recovery from the collected droplets. We also observed that the collected droplets begin to shrink after the collection device is disconnected from the sorting device and that shrinkage is less severe when larger numbers of droplets are collected. We assessed whether viability is affected by the sorting process itself by running cell-containing droplets through the device either into the waste channel (electrode turned off) or into the collection channel (electrode turned on), collecting the droplets in Eppendorf tubes and assessing the cell viability. We found no significant difference between the droplets that had not been reinjected, the waste channel droplets, and the positive channel droplets (1.7×10^4^ ± 3.4×10^3^, 2.0×10^4^ ± 3.2×10^3^, and 1.6×10^4^ ± 88 CFU/μL droplets (SD, n = 2), respectively). We, therefore, conclude that droplet shrinkage is likely to cause incomplete cell recovery. Subsequently, as an alternative and direct way of examining sorting efficiency, we labeled the high secretor JCL260 Δ*lysA* pSA69 with mCherry by transformation with ampicillin-resistant pBT-proD-mCherry. To enable cultivation in ampicillin, lower secretor JCL16 Δ*lysA* was transformed with empty pTGD plasmid and K12 *DilvD DgalK::cfp-bla* pSAS31 (expressing both CFP and mNeongreen) was employed as the sensor strain. Thus, droplets could be sorted for green fluorescence and accuracy could be assessed based on whether droplets contained mCherry-expressing cells.

We prepared droplets with a mixture of JCL260 Δ*lysA* pSA69 and JCL16 Δ*lysA* secretor strains, at an initial ratio of 1:1,000, on 1.0 g/L norvaline. In 30 min of sorting at a droplet reinjection rate of 2 μL/min (~400,000 droplets total, or ~40,000 containing a secretor strain), we retrieved five droplets in the collection device. All five contained substantial numbers of green fluorescent cells, while four contained red fluorescent cells, indicative of the high secretor (true positive) (Fig. S13). This demonstrates the accuracy of the sorting system at high throughput (i.e., it is possible to sort a large number of droplets and isolate only those containing a high number of green cells) and the low biological false positive rate.

## 5. Conclusion

In this study, we have developed the SnoCAP framework for converting inconspicuous production phenotypes into growth phenotypes, facilitating high-throughput screening of production strain libraries via compartmentalization of cross-feeding production and sensor strains. The use of metabolite analogs makes the assay dynamic range highly tunable without requiring any genetic modifications. Assay timing can also be adjusted by kick-starting the culture with low-concentrations of the cross-fed metabolites. Since assay readout is cell density or cell fluorescence, no costly assay reagents are necessary. We have shown the utility of this method for screening strain libraries and identified a strain that overproduced 2-KIV without a plasmid, which is desirable since plasmids require antibiotics for maintenance, and even then may still be lost under non-ideal conditions (Minty et al. 2013).

We have demonstrated three implementation formats, each with its own advantages and limitations. The microtiter plate assay, although requiring the most space, avoids the issue of single cell variability and therefore provides the highest accuracy, with the tightest agreement of replicates, both between and within experiments. The agar plate format reaches high throughputs (up to ~10^5^ assays per square meter of plate surface) and does not require specialized equipment. The high cell density possible in a colony enables a large number of doublings and hence a large degree of amplification of differences in production phenotype. The microdroplet format achieves ultrahigh throughputs (~10^6^ droplets/h, corresponding to ~10^5^ assays/h, or ~10^6^ assays/day, when a secretor loading of ~0.1/droplet is used) that can drastically reduce screening time and enable screening of libraries that are orders of magnitude larger. The small culture volume also significantly shortens the requisite incubation periods. As with most high-throughput single cell tools, single cell variability decreases the precision of the agar plate and microdroplet formats. In the microdroplet assay, there is additional variability between replicates because the cells are encapsulated according to the Poisson distribution, and each droplet does not start with an identical number of sensor cells. A more sophisticated incubation setup for improved aeration (e.g., in the work of Mahler et al. (2015)) may also improve the homogeneity of the conditions experienced by each droplet. Advances in absorbance-activated droplet sorting (such as that reported by Gielen et al. (2016)) may also enable application of the droplet format to systems in which the strains are not easily made fluorescent.

In our screening of a chemically mutagenized library using the agar plate format of SnoCAP, we identified a strain with superior production level but compromised growth. The higher throughput capability of the microdroplet assay format would likely enable identification of a larger number of high producers, which could then be subjected to a second round of screening or selection for high monoculture growth rate to search for strains that are not growth-compromised.

We expect that the SnoCAP screening framework can be applied to a wide variety of industrially relevant target molecules. The primary requirements for application of SnoCAP are that the molecule whose production level is being screened for can be extracellularly secreted and that an auxotrophic strain exists or can be constructed. By a push-pull strategy, it can also be extended to secondary metabolites that are several steps removed from a primary metabolite. We also expect that this technology can be applied to various other microbial species. Synthetic cross-feeding consortia have been examined in yeast (Shou, Ram, and Vilar 2007), and *E. coli* has shown the ability to cross-feed with other species, such as *Acinetobacter baylyi* (Pande et al. 2015) or *Salmonella* species (Harcombe et al. 2018; Fildes 1956). The strategy could also be extended to screening for overproduction of other compounds of interest that a strain with a diverse metabolism, such as *Pseudomonas putida*, can utilize as a carbon source but the secretor strain cannot. In this case, a carbon source would be supplied to the secretor that the sensor cannot utilize.

## Acknowledgments

We thank Prof. James Liao (UCLA/Academia Sinica) for sharing strains JCL16, JCL260 and plasmids pSA65, pSA69, and Prof. Keith Tyo (Northwestern University) for plasmid pTGD. We are also grateful to Brian Johnson for 3D printing Petri dish holders for the agar plate scanning assays, Scott Scholz for helpful discussions and for providing the pSAS31 plasmid, James Windak in the University of Michigan Department of Chemistry for technical assistance in the measurement of 2-KIV, and Profs. Lola Eniola, Mark Burns, Jinsang Kim, and Allen Liu (University of Michigan) for generously allowing us to use equipment in their laboratories. This work was supported by the USDA AFRI NIFA Fellowships Grant Program (Grant no. 2016-67011-24725).

## References

J. Abatemarco, M.F. Sarhan, J.M. Wagner, J.L. Lin, L. Liu, W. Hassouneh, S.F. Yuan, H.S. Alper, and A.R. Abate. RNA-aptamers-in-droplets (RAPID) high-throughput screening for secretory phenotypes. Nature Communications, 8 (2017): 332.

S. Atsumi, T. Hanai, and J.C. Liao. Non-fermentative pathways for synthesis of branched-chain higher alcohols as biofuels. Nature, 451 (2008): 86.

S. Atsumi, T.-Y. Wu, E.-M. Eckl, S.D. Hawkins, T. Buelter, and J.C. Liao. Engineering the isobutanol biosynthetic pathway in *Escherichia coli* by comparison of three aldehyde reductase/alcohol dehydrogenase genes. Applied Microbiology and Biotechnology, 85 (2010): 651–57.

J.-C. Baret, O.J. Miller, V. Taly, M. Ryckelynck, A. El-Harrak, L. Frenz, C. Rick, M.L. Samuels, J.B. Hutchison, and J.J. Agresti. Fluorescence-activated droplet sorting (FADS): efficient microfluidic cell sorting based on enzymatic activity. Lab on a Chip, 9 (2009): 1850–58.

G.W. Beadle, and E.L. Tatum. Genetic Control of Biochemical Reactions in Neurospora. Proceedings of the National Academy of Sciences of the United States of America, 27 (1941): 499–506.

F. Bertels, H. Merker, and C. Kost. Design and characterization of auxotrophy-based amino acid biosensors. PLoS One, 7 (2012): e41349.

P. R. Burkholder. Determination of vitamin B12 with a mutant strain of *Escherichia coli*. Science, 114 (1951): 459–60.

J.G.R. Cardoso, A.A. Zeidan, K. Jensen, N. Sonnenschein, A.R. Neves, and M.J. Herrgard. MARSI: metabolite analogues for rational strain improvement. Bioinformatics, 34 (2018): 2319–21.

P.A. Carr, H.H. Wang, B. Sterling, F.J. Isaacs, M.J. Lajoie, G. Xu, G.M. Church, and J.M. Jacobson. Enhanced multiplex genome engineering through co-operative oligonucleotide co-selection. Nucleic Acids Research, 40 (2012): e132–e32.

V. Chalova, W. Kim, C. Woodward, and S. Ricke. Quantification of total and bioavailable lysine in feed protein sources by a whole-cell green fluorescent protein growth-based *Escherichia coli* biosensor. Applied Microbiology and Biotechnology, 76 (2007): 91–99.

V.I. Chalova, S.A. Sirsat, C.A. O’Bryan, P.G. Crandall, and S.C. Ricke. *Escherichia coli*, an intestinal microorganism, as a biosensor for quantification of amino acid bioavailability. Sensors, 9 (2009): 7038–57.

M.T. Chung, D. Núñez, D. Cai, and K. Kurabayashi. Deterministic droplet-based coencapsulation and pairing of microparticles via active sorting and downstream merging. Lab on a Chip, 17 (2017): 3664–71.

J.A. Dietrich, A.E. McKee, and J.D. Keasling. High-throughput metabolic engineering: advances in small-molecule screening and selection. Annual Review of Biochemistry, 79 (2010): 563–90.

M. Fang, T. Wang, C. Zhang, J. Bai, X. Zheng, X. Zhao, C. Lou, and X.-H. Xing. Intermediate-sensor assisted push-pull strategy and its application in heterologous deoxyviolacein production in *Escherichia coli*. Metabolic Engineering, 33 (2016): 41–51.

P. Fildes. Production of tryptophan by *Salmonella typhi* and *Escherichia coli*. Microbiology, 15 (1956): 636–42.

F. Gielen, R. Hours, S. Emond, M. Fischlechner, U. Schell, and F. Hollfelder. Ultrahigh-throughput-directed enzyme evolution by absorbance-activated droplet sorting (AADS). Proceedings of the National Academy of Sciences, 113 (2016): E7383–E89.

A. D. Haimovich, P. Muir, and F.J. Isaacs. Genomes by design. Nature Reviews Genetics, 16 (2015): 501–16.

W.R. Harcombe, J.M. Chacón, E.M. Adamowicz, L.M. Chubiz, and C.J. Marx. Evolution of bidirectional costly mutualism from byproduct consumption. Proceedings of the National Academy of Sciences, 115 (2018): 12000–04.

M. Huang, Y. Bai, S.L. Sjostrom, B.M. Hallström, Z. Liu, D. Petranovic, M. Uhlén, H.N. Joensson, H. Andersson-Svahn, and J. Nielsen. Microfluidic screening and whole-genome sequencing identifies mutations associated with improved protein secretion by yeast. Proceedings of the National Academy of Sciences, 112 (2015): E4689–E96.

O. Karlström. Methods for the production of mutants suitable as amino acid fermentation organisms. Biotechnology and Bioengineering, 7 (1965): 245–68.

A. Kerner, J. Park, A. Williams, and X.N. Lin. A programmable *Escherichia coli* consortium via tunable symbiosis. PLoS One, 7 (2012): e34032.

M. I. Kim, T.J. Park, N.S. Heo, M.-A. Woo, D. Cho, S.Y. Lee, and H.G. Park. Cell-based method utilizing fluorescent *Escherichia coli* auxotrophs for quantification of multiple amino acids. Analytical Chemistry, 86 (2014): 2489–96.

I. Levin-Reisman, O. Fridman, and N.Q. Balaban. ScanLag: High-throughput Quantification of Colony Growth and Lag Time. Journal of Visual Experiments, 89 (2014)

I. Levin-Reisman, O. Gefen, O. Fridman, I. Ronin, D. Shwa, H. Sheftel, and N.Q. Balaban. Automated imaging with ScanLag reveals previously undetectable bacterial growth phenotypes. Nature Methods, 7 (2010): 737–39.

M. Li, M. van Zee, C.T. Riche, B. Tofig, S.D. Gallaher, S.S. Merchant, R. Damoiseaux, K. Goda, and D. Di Carlo. A gelatin microdroplet platform for high-throughput sorting of hyperproducing single-cell-derived microalgal clones. Small (2018): 1803315.

J.-L. Lin, J.M. Wagner, and H.S. Alper. Enabling tools for high-throughput detection of metabolites: Metabolic engineering and directed evolution applications. Biotechnology Advances, 35 (2017): 950–70.

S.N. Lindner, L.C. Ramirez, J.L. Krüsemann, O. Yishai, S. Belkhelfa, H. He, M. Bouzon, V. Döring, and A. Bar-Even. NADPH-Auxotrophic E. coli: A Sensor Strain for Testing in Vivo Regeneration of NADPH. ACS Synthetic Biology, 7 (2018): 2742–49.

G. Lopez, and J.C. Anderson. Synthetic Auxotrophs with Ligand-Dependent Essential Genes for a BL21(DE3) Biosafety Strain. ACS Synthetic Biology, 4 (2015): 1279–86.

L. Mahler, M. Tovar, T. Weber, S. Brandes, M.M. Rudolph, J. Ehgartner, T. Mayr, M.T. Figge, M. Roth, and E. Zang. Enhanced and homogeneous oxygen availability during incubation of microfluidic droplets. RSC Advances, 5 (2015): 101871–78.

M.T. Mee, J.J. Collins, G.M. Church, and H.H. Wang. Syntrophic exchange in synthetic microbial communities. Proceedings of the National Academy of Sciences, 111 (2014): E2149–E56.

J.J. Minty, M.E. Singer, S.A. Scholz, C.-H. Bae, J.-H. Ahn, C.E. Foster, J.C. Liao, and X.N. Lin. Design and characterization of synthetic fungal-bacterial consortia for direct production of isobutanol from cellulosic biomass. Proceedings of the National Academy of Sciences, 110 (2013): 14592–97.

X. Niu, F. Gielen, J.B. Edel, and A.J. deMello. A microdroplet dilutor for high-throughput screening. Nature Chemistry, 3 (2011): 437.

V. Nurmikko. Biochemical Factors Affecting Symbiosis Among Bacteria. Experientia, 12 (1956): 245–84.

S. Pande, S. Shitut, L. Freund, M. Westermann, F. Bertels, C. Colesie, I.B. Bischofs, and C. Kost. Metabolic cross-feeding via intercellular nanotubes among bacteria. Nature Communications, 6 (2015): 6238.

J. Park, A. Kerner, M.A. Burns, and X.N. Lin. Microdroplet-enabled highly parallel cocultivation of microbial communities. PloS One, 6 (2011): e17019.

M. Park, S.-L. Tsai, and W. Chen. Microbial biosensors: Engineered microorganisms as the sensing machinery. Sensors, 13 (2013): 5777–95.

J. Payne, G. Bell, and C. Higgins. The use of an *Escherichia coli* Lys-auxotroph to assay nutritionally available lysine in biological materials. Journal of Applied Microbiology, 42 (1977): 165–77.

B. F. Pfleger, D.J. Pitera, J.D. Newman, V.J. Martin, and J.D. Keasling. Microbial sensors for small molecules: Development of a mevalonate biosensor. Metabolic Engineering, 9 (2007): 30–38.

S.A. Scholz, R. Diao, M.B. Wolfe, E.M. Fivenson, X.N. Lin, and P.L. Freddolino. High-resolution mapping of a standardized transcriptional reporter reveals dedicated high and low transcription domains in the *Escherichia coli* chromosome. Revision submitted to Cell Systems (2018).

W. Shou, S. Ram, and J.M. Vilar. Synthetic cooperation in engineered yeast populations. Proceedings of the National Academy of Sciences, 104 (2007): 1877–82.

S. Siedler, N.K. Khatri, A. Zsohár, I. Kjærbølling, M. Vogt, P. Hammar, C.F. Nielsen, J. Marienhagen, M.O.A. Sommer, and H.N. Joensson. Development of a bacterial biosensor for rapid screening of yeast p-coumaric acid production. ACS Synthetic Biology, 6 (2017): 1860–69.

K.M. Smith, and J.C. Liao. An evolutionary strategy for isobutanol production strain development in *Escherichia coli*. Metabolic Engineering, 13 (2011): 674–81.

N. Tepper, and T. Shlomi. Computational design of auxotrophy-dependent microbial biosensors for combinatorial metabolic engineering experiments. PLOS One, 6 (2011): e16274.

K.E.J. Tyo, P.K. Ajikumar, and G. Stephanopoulos. Stabilized gene duplication enables longterm selection-free heterologous pathway expression. Nature Biotechnology, 27 (2009): 760–65.

F. Valle, and A. Berry. Metabolic Engineering of Escherichia coli for the Production of Aromatic Compounds. In: Metabolic Engineering, Lee, S. Y., and Papoutsakis, E. T. Eds. Marcel Dekker, Inc., New York, 79 (1999). (Taylor & Francis).

J.M. Wagner, L. Liu, S.-F. Yuan, M.V. Venkataraman, A.R. Abate, and H.S. Alper. A comparative analysis of single cell and droplet-based FACS for improving production phenotypes: riboflavin overproduction in *Yarrowia lipolytica*. Metabolic Engineering, 47 (2018): 346–56.

B. L. Wang, A. Ghaderi, H. Zhou, J. Agresti, D.A. Weitz, G.R. Fink, and G. Stephanopoulos. Microfluidic high-throughput culturing of single cells for selection based on extracellular metabolite production or consumption. Nature Biotechnology, 32 (2014): 473.

K.C. Winkler, A.W. van Doorn, and A.F. Royers. Symbiosis of tryptophan-deficient mutants of *E*. coli B. Recl. Trav. Chim. Pays-Bas, 71 (1952): 5–14.

E.H. Wintermute, and P.A. Silver. Emergent cooperation in microbial metabolism. Molecular Systems Biology, 6 (2010): 407–07.

J.-H. Zhang, T.D. Chung, and K.R. Oldenburg. A simple statistical parameter for use in evaluation and validation of high throughput screening assays. Journal of Biomolecular Screening, 4 (1999): 67–73.

X. Zhang, and J.L. Reed. Adaptive evolution of synthetic cooperating communities improves growth performance. PloS One, 9 (2014): e108297.

Z.-J. Zhao, C. Zou, Y.-X. Zhu, J. Dai, S. Chen, D. Wu, J. Wu, and J. Chen. Development of L-tryptophan production strains by defined genetic modification in *Escherichia coli*. Journal of Industrial Microbiology & Biotechnology, 38 (2011): 1921–29.

J. Zheng, and Z. Jia. Structure of the bifunctional isocitrate dehydrogenase kinase/phosphatase. Nature, 465 (2010): 961.

