## Supplementary Information for "Syntrophic co-culture amplification of production phenotype for high-throughput screening of microbial strain libraries"

**This PDF file includes:**

Figs. S1 to S13 and Supplementary Notes 1 & 2……………….……………..……..…2
Tables S1 and S2……………………………………………………..……..…...…17
Supplementary Methods………………...…………..……….....………………..…..21
Caption for Dataset S1……………………….……...……..…………….……..…29
Captions for Movies S1 and S2……………...……………..……………..………..…29
Supplementary References……………………………………..…………….……..…30

**Other supplementary materials for this manuscript include the following:**

Dataset S1
Movies S1 and S2

### Supplementary Figures and Notes

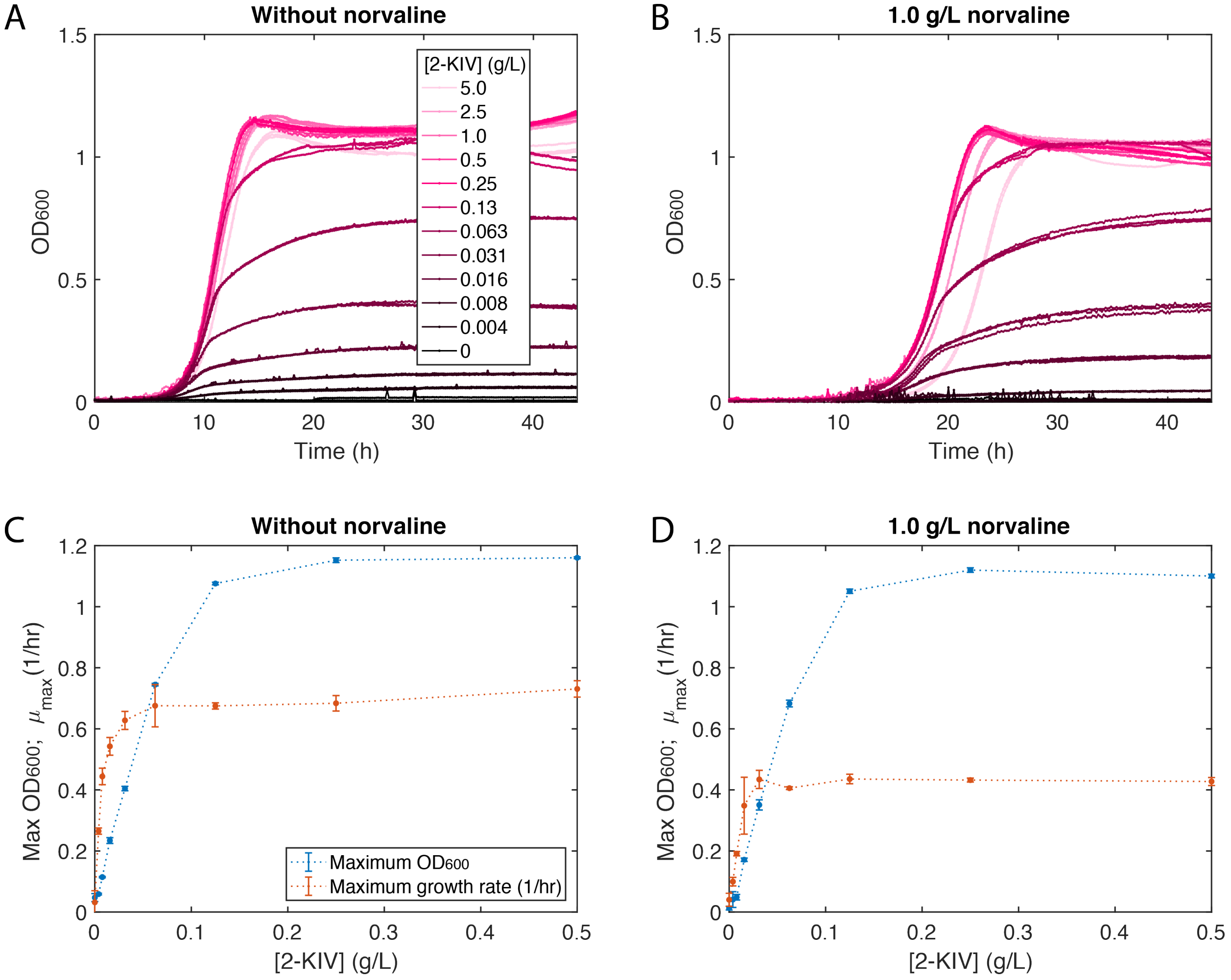

**Figure S1:** Sensitivity of K12 Δ*ilvD* sensor growth characteristics to 2-KIV concentration in monoculture. Growth profiles without (A) and with 1.0 g/L (B) norvaline. The medium contains an excess of isoleucine. Three replicate wells are plotted in the same color for each 2-KIV concentration. (C, D) Maximum $OD_{600}$ and growth rate from the growth profiles in (A, B).

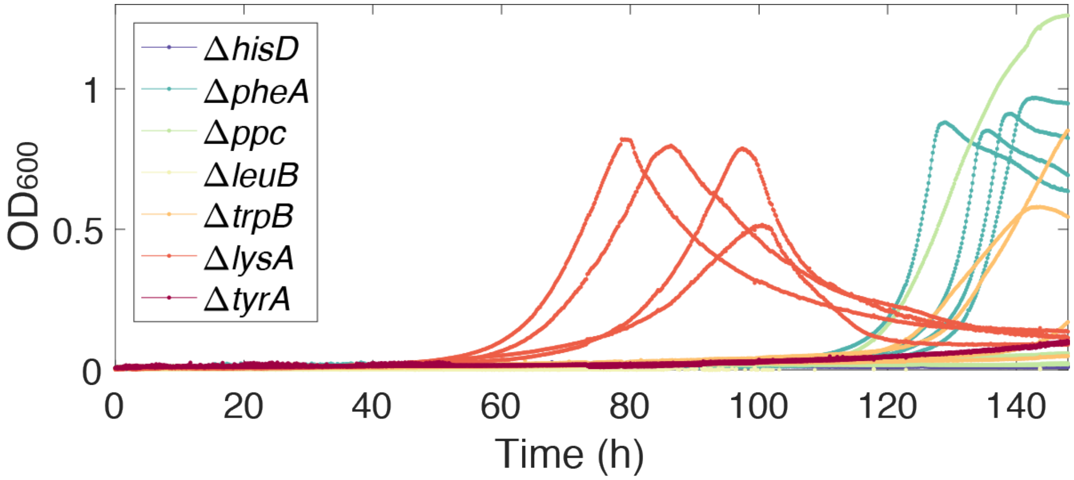

**Figure S2:** Co-culture growth profiles of various Keio strains with K12 Δ*ilvD*. Biological replicates are plotted in the same color (n = 4). Legend denotes the gene knockout of the Keio strain.

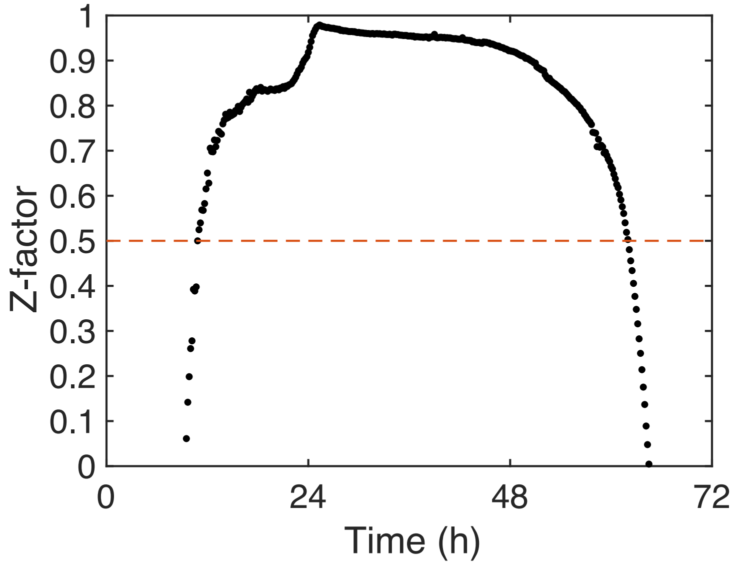

**Figure S3:** Z-factors over time calculated from the $OD_{600}$ readings of co-cultures containing secretor strain JCL16 Δ*lysA* as negative control and JCL260 Δ*lysA* pSA69 as positive control (8 replicates each; medium containing 0.1 mM IPTG and no norvaline). The Z-factor is defined as $Z = 1 - \frac{3\sigma_p+3\sigma_n}{|\mu_p-\mu_n|}$, where $\mu$ and $\sigma$ are the mean and standard deviation of the positive ($p$) and negative ($n$) controls.

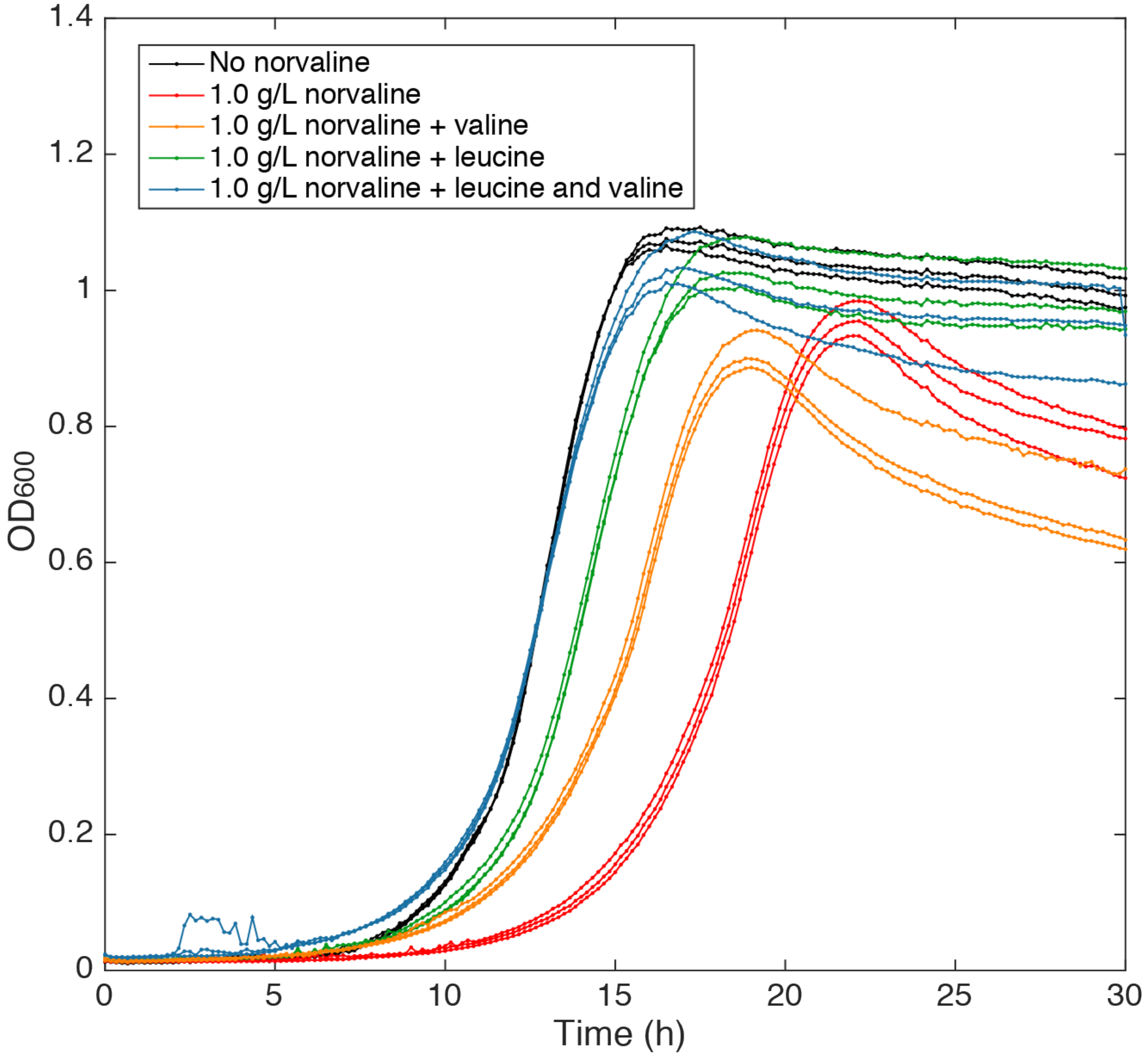

**Figure S4:** Growth profiles of JCL16 Δ*lysA* supplemented with 3 mM lysine and 3 mM isoleucine with and without 1.0 g/L norvaline, 3 mM valine, and 3 mM leucine. Three biological replicates of each culture are plotted in the same color.

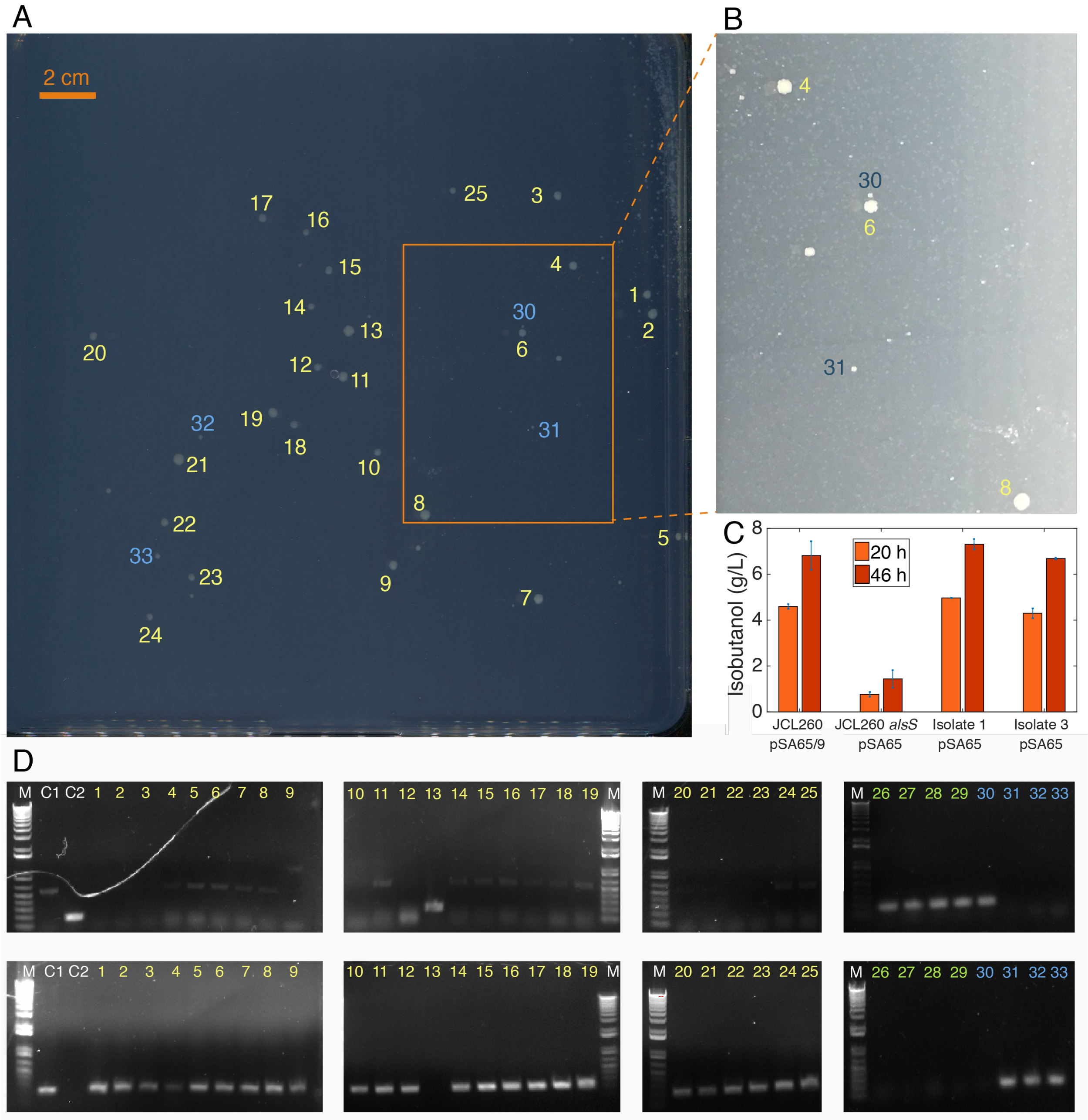

**Figure S5:** Model library on agar plate. (A) 0.6% library plate after incubation at 37 °C for 7 days, imaged from below on photo scanner. Colonies that were further examined are numbered. (B) A close-up of the plate, photographed from above. Many small colonies are visible surrounding the large colonies. (C) Production levels of secretor strain isolated from colonies 1 and 3 in the screen after transformation with pSA65, in comparison with JCL260 Δ*lysA* pSA65/9 and JCL260 Δ*lysA alsS* pSA65. Error bars represent standard deviation between two biological replicates. (D) Verification that large colonies in the model library are JCL260 Δ*lysA* pSA69 and not JCL260 Δ*lysA alsS*. Gel electrophoresis of PCR reactions with either primers alsS_int_chk_front_for and alsS_int_chk_front_rev (top) or alsS_int_chk_front_for and alsS_int_chk_back_rev (bottom). M is Invitrogen 1 Kb plus ladder, C1 is JCL260 Δ*lysA* pSA69, C2 is JCL260 Δ*lysA alsS.* 1-25 are secretor strain colonies isolated from the 25 largest mixed colonies. 26-29 are secretor strain isolated from four small colonies from the model library plate. 30-33 are intermediate-sized colonies. Numbers correspond to the numbering shown on Fig. S5A, except for the small colonies which are not visible in that image.

**Supplementary Note 1:** Results of screening of model libraries on agar plates.

*0.6% library:*

A square plate (shown in Fig. S5) was spread with 102 ± 25 LB-CFU of JCL260 Δ*lysA* pSA69 and 18,200 ± 500 of JCL260 Δ*lysA alsS* (based on plating the inocula on LB plates; SD, n = 2). After 7 days, 25 large colonies were apparent and are numbered in yellow in Fig. S5. 17 intermediate-sized colonies were also observed. The four that were further investigated are numbered in blue (Fig. S5A,B).

Numbered colonies, as well as 4 small colonies, were streaked on LB plates with tetracycline and kanamycin to isolate secretor strain. A single colony from each of these plates was assayed by PCR using primer sets alsS_int_chk_front_for/alsS_int_chk_front_rev, which produce a short band only if *alsS* is integrated, and alsS_int_chk_front_for/alsS_int_chk_back_rev, which produce a short band only if *alsS* is not integrated (Fig. S5D). 24 of the 25 large colonies were identified as JCL260 Δ*lysA* pSA69.

Colony 13, the only large colony which was identified as containing JCL260 Δ*lysA alsS*, was a particularly large colony, so we suspected it could have merged with surrounding small colonies, leading to a mixture of secretor strains. We repeated the streak out of Colony 13 on an LB tetracycline plate, picking cells from the very center of the colony. When we repeated the PCR assay on 10 colonies from this streak, we found that 9 of 10 were JCL260 Δ*lysA* pSA69 and only one was JCL260 Δ*lysA alsS*.

The four small colonies that were picked were all identified as JCL260 Δ*lysA alsS*, as expected. Of the four intermediate-sized colonies that were investigated, three were JCL260 Δ*lysA* pSA69 and one (colony 30) was JCL260 Δ*lysA alsS.* Colony 30 is quite close to colony 6, so it is possible that diffusion between colonies in close proximity is occurring, which should be taken into consideration when selecting colonies.

*0.1% library:*

A 0.1% model library was also tested. We plated 9.1 ± 1.6 LB-CFU of JCL260 Δ*lysA* pSA69 and 78,500 ± 70 LB-CFU of JCL260 Δ*lysA alsS* on each of two square plates. After six days incubation at 37 $^{o}$C, one plate had developed one large colony and the other developed two. All three large colonies were found to contain JCL260 Δ*lysA* pSA69 as the secretor strain.

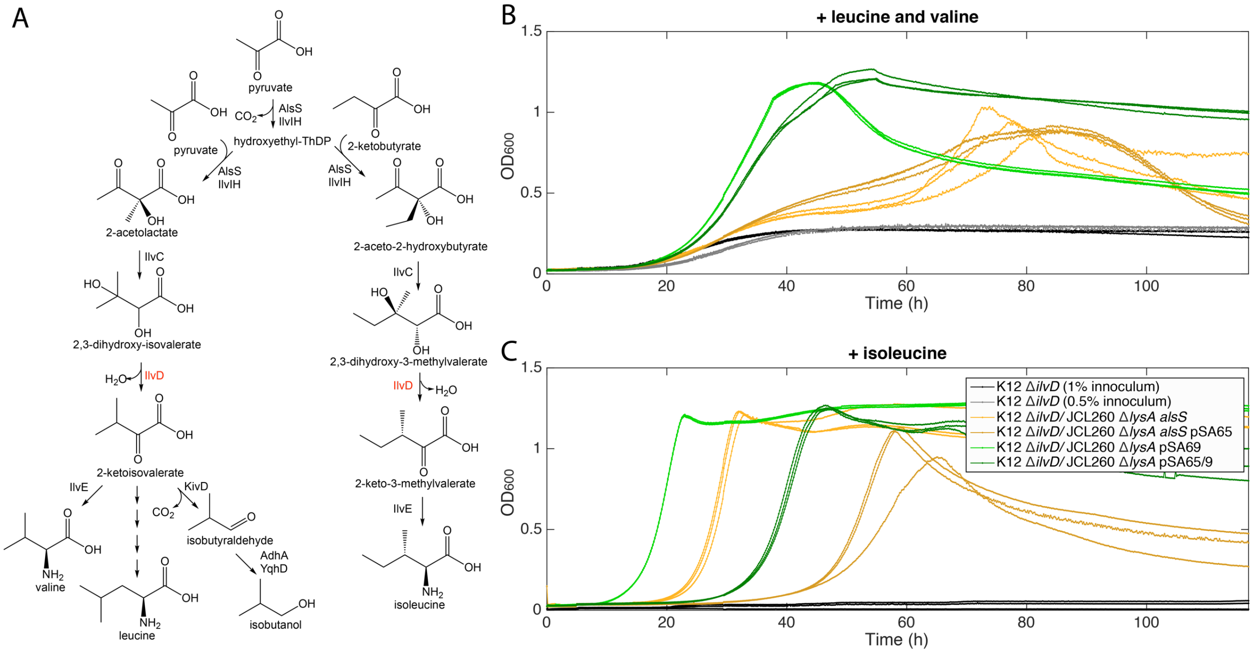

**Figure S6:** Investigation of the cause of the reduced co-culture growth in strains with increased alpha-ketoisovalerate decarboxylase and alcohol dehydrogenase activity. (A) Branched-chain amino acid and isobutanol pathways. Co-culture and sensor strain monoculture growth profiles in liquid medium supplemented with either 3 mM of both leucine and valine (B) or 3 mM isoleucine (C).

**Supplementary Note 2:** Validation of cause of decreased co-culture growth with plasmid-carrying production strain.

Because the addition of pSA65 adds metabolic burden and a toxic product, it was not clear how much of the decrease in growth was indeed due to 2-KIV being channeled away from cross-feeding. We, therefore, tested these co-cultures under the condition of excess leucine and valine instead of isoleucine. This condition leads to a co-culture that must cross-feed isoleucine or intermediates for the production of isoleucine, and we do not expect channeling of 2-KIV to affect the co-culture fitness (Fig. S6A). If defects in growth were due mainly to factors other than

2-KIV channeling, such as a metabolic burden or product toxicity, then we would expect them to be observed under the leucine/valine supplementation as well. Under these conditions, we observed some basal growth of the sensor strain monoculture, but the co-cultures have significantly more growth. For both secretor strains tested (JCL260 Δ*lysA alsS* and JCL260 Δ*lysA* pSA69), the version carrying pSA65 has reduced growth compared to that without pSA65 under the isoleucine-supplemented condition (Fig. S6C) and has similar growth to that without pSA65 under the leucine/valine-supplemented condition (Fig. S6B). These results support the prediction that the co-growth characteristics will be determined primarily by the production level of the cross-fed molecules

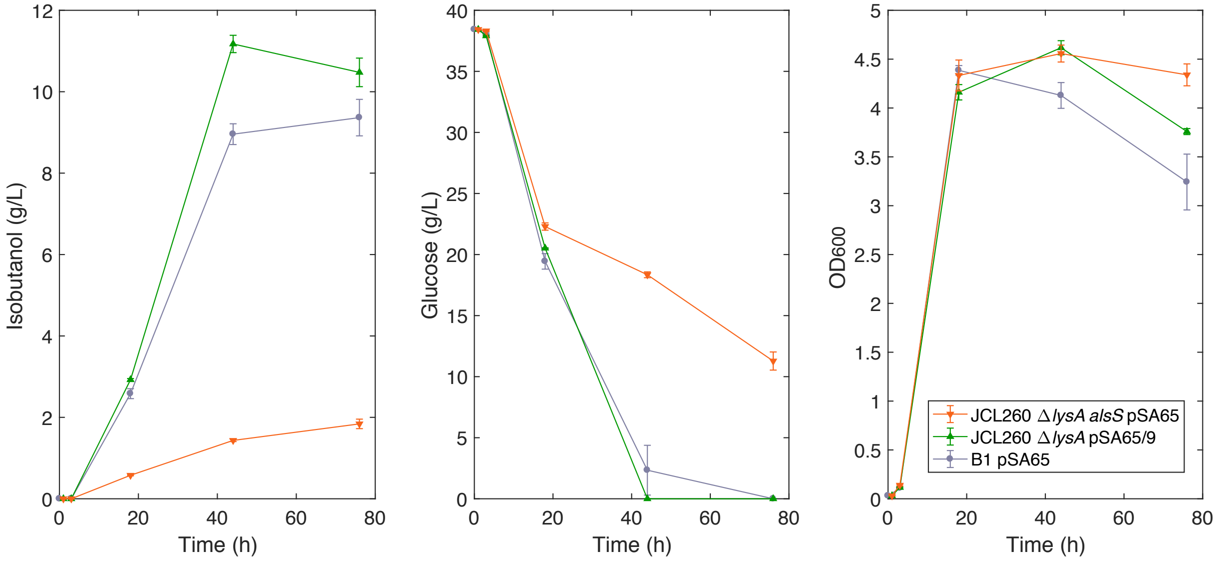

**Figure S7:** Production culture performance of mutagenesis library isolate B1 compared to parent strain JCL260 Δ*lysA alsS* and high-producing strain JCL260 Δ*lysA* pSA69. Error bars represent standard deviations of biological replicates (n = 3).

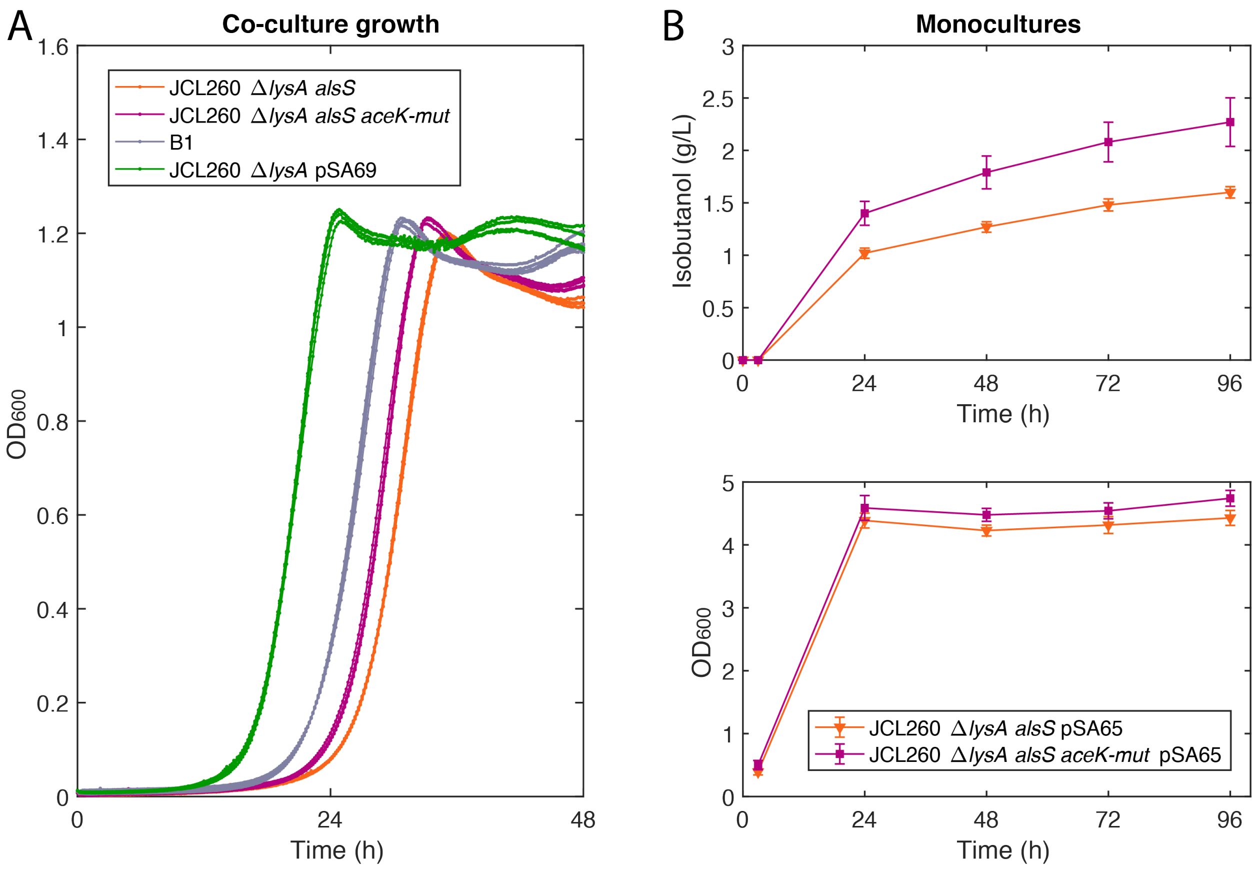

**Figure S8**: *aceK* mutant strain characterization. (A) Growth profiles of secretor strains co-cultured with K12 Δ*ilvD* sensor strain in a 96-well microplate with 0.1 g/L norvaline (n = 4, replicates plotted in the same color). (B) Monoculture isobutanol production and growth profiles of *aceK* mutant strain and parental strain after transformation with pSA65. Error bars represent the standard deviation of biological replicates (n = 10).

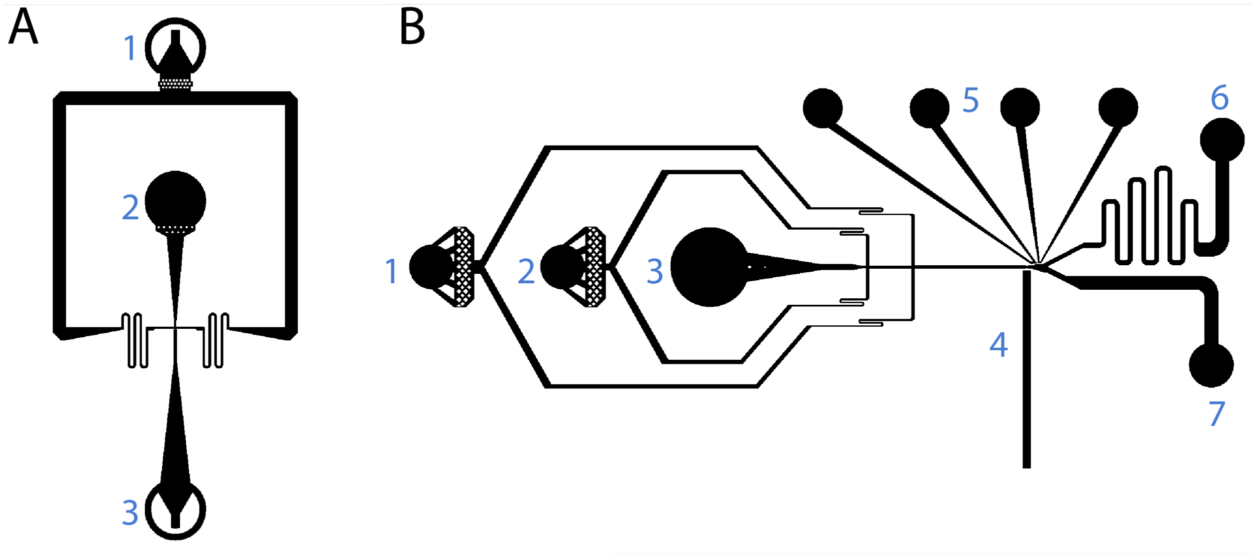

**Figure S9:** Microfluidic devices. (A) Flow-focusing droplet generation device used to encapsulate cells. Labeled components: (1) oil inlet, (2) cell suspension inlet, (3) outlet. (B) Droplet sorting device. Labeled components: (1, 2) spacing oil inlets, (3) droplet reinjection port, (4) optical fiber channel, (5) electrodes, (6) desired droplet outlet, (7) undesired droplet outlet.

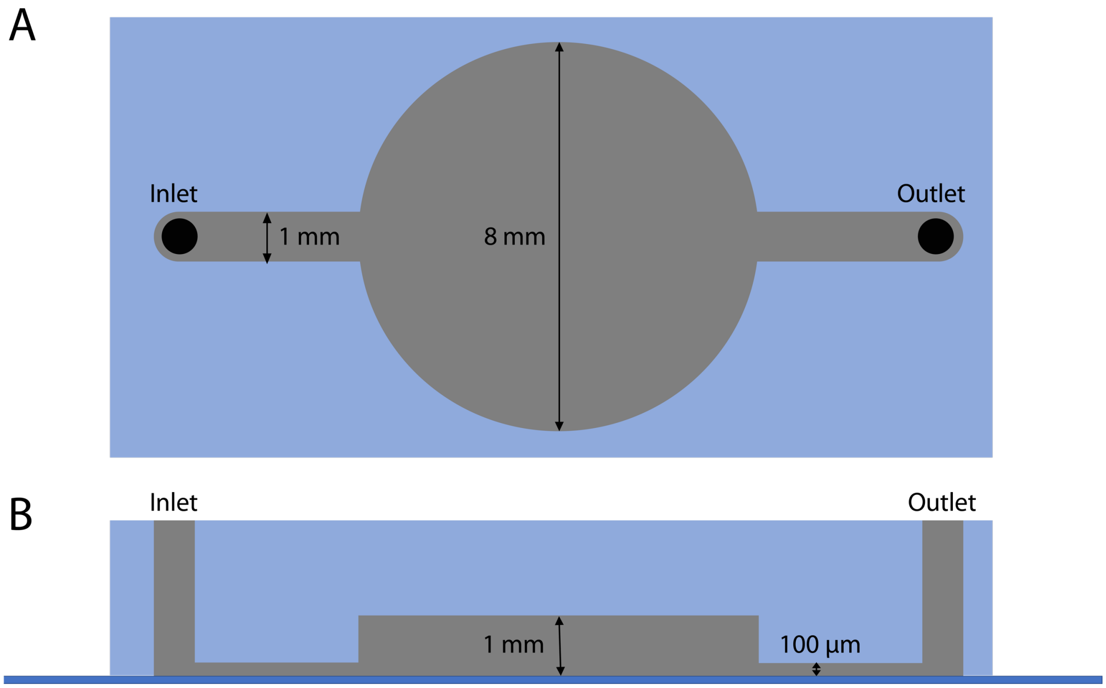

**Figure S10:** Droplet collection device. Top (A) and side (B) views.

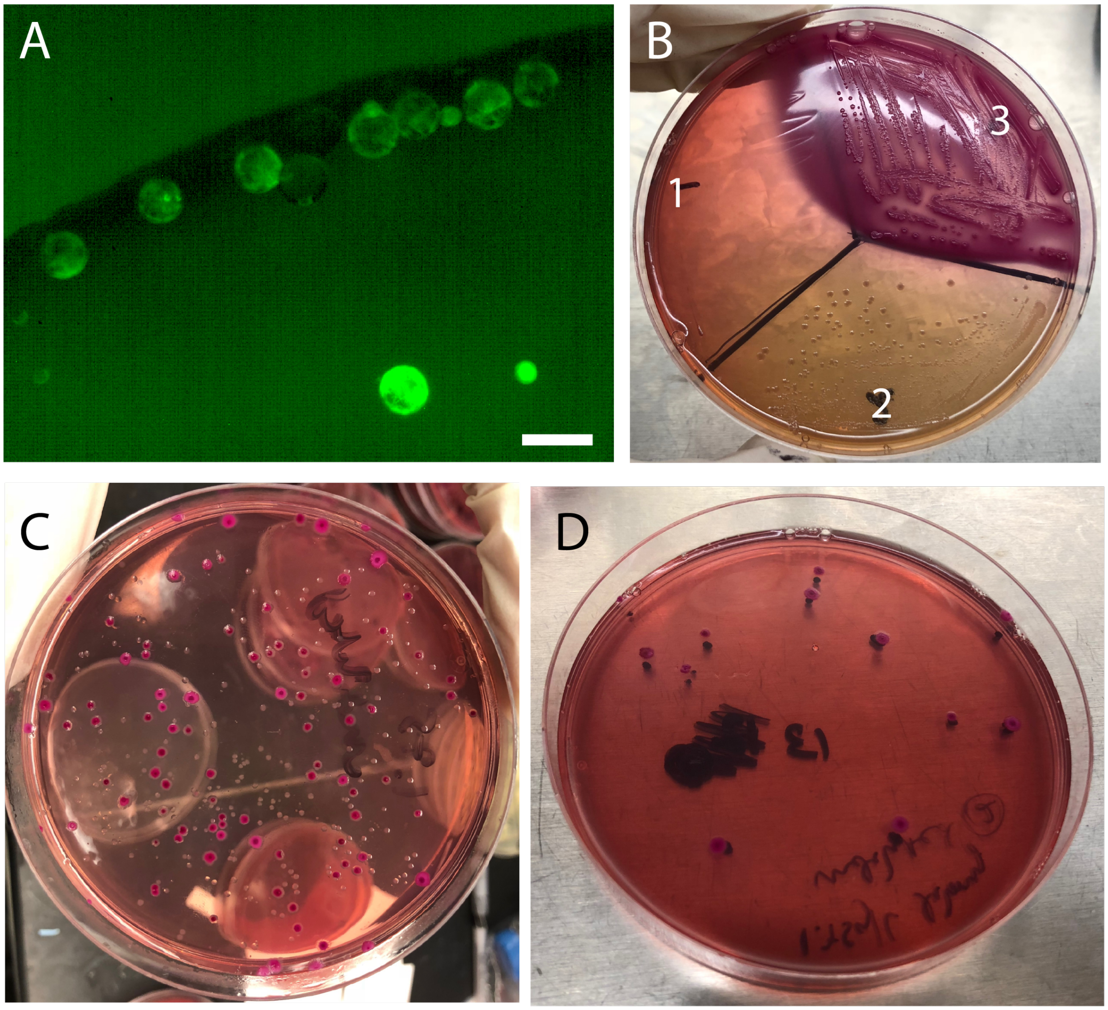

**Figure S11:** Retrieval of viable cells after droplet sorting. (A) Droplets collected after 25 min sorting of droplets encapsulating a mixture of JCL260 Δ*lysA alsS* and JCL260 Δ*lysA* pSA69 secretor strains at a 100:1 ratio. Scale bar: 100 µm. (B) Strain growth on MacConkey agar plates containing tetracycline. K12 Δ*ilvD* pSAS31 (1) does not grow. JCL260 Δ*lysA alsS* Δ*galK* (2) produces white colonies, and JCL260 Δ*lysA* pSA69 (3) produces purple colonies. (C) Colonies from plating unsorted droplets. (D) Colonies from plating the collected droplets that are pictured in (A).

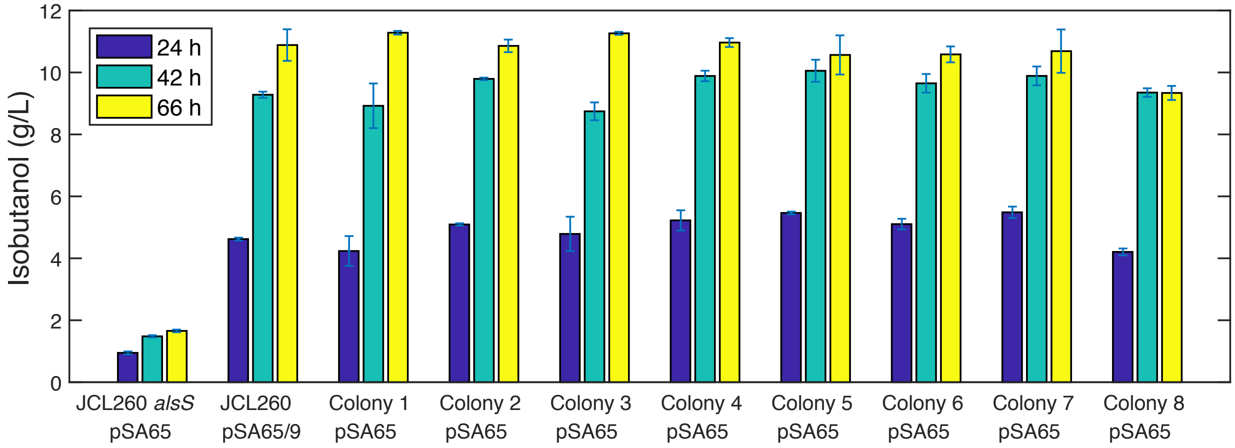

**Figure S12:** Isobutanol production of cells from eight randomly selected purple colonies isolated from sorting of model libraries and transformed with pSA65, compared to JCL260 Δ*lysA alsS* pSA65 and JCL260 Δ*lysA* pSA65/9. Error bars represent standard deviation of two biological replicates.

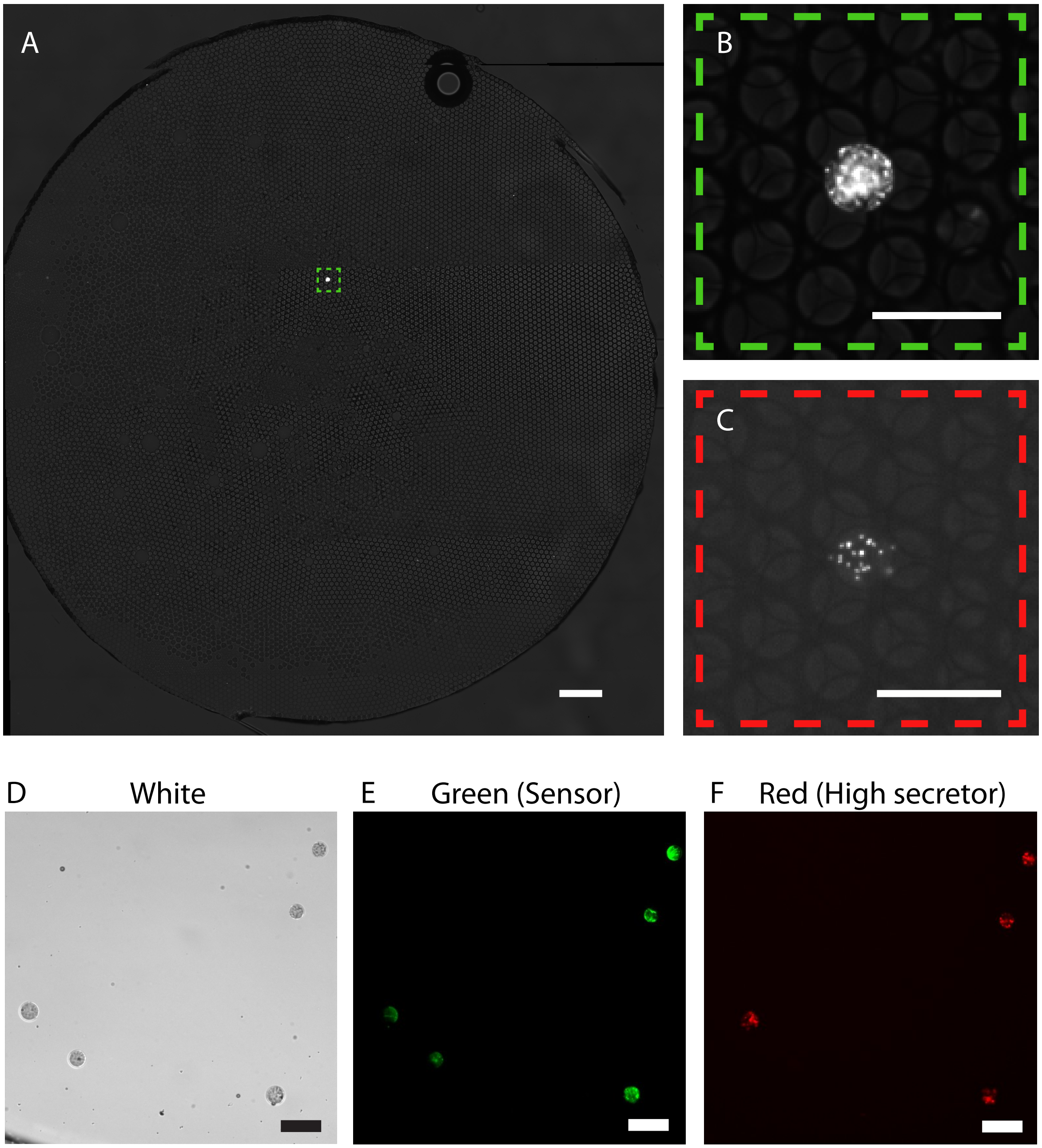

**Figure S13:** Droplets before and after fluorescence-activated droplet sorting (FADS). (A) ~90 pL droplets were generated containing sensor strain at a cell loading of ~5 sensor cells/droplet and secretor strains JCL16 Δ*lysA* pTGD and JCL260 Δ*lysA* pSA69/pBT1-proD-mCherry, mixed at a ratio of 1,000:1, with a total loading of ~0.1 secretor cell/droplet. After 36 h incubation, a sample of these droplets was examined. One droplet with substantial sensor cell growth is visible from among ~5*$10^4$ droplets (A, B). This droplet was identified as containing JCL260 Δ*lysA* pSA69/pBT1-proD-mCherry secretor strain by observation of cells expressing mCherry (C). After 30 min FADS of these droplets, five droplets were collected (D-F). All five showed growth of the sensor strain as determined by cells with green fluorescence (E) and four contained cells with red fluorescence, indicative of JCL260 Δ*lysA* pSA69/pBT1-proD-mCherry (F). Scale bars: (A) 500 μm, (B-F) 100 μm.

#### Supplementary Tables

##### Table S1: Strains and plasmids

| Strain | Relevant genotype | Reference |
|---|---|---|
| JCL16 | BW25113/F′ [*traD36*, *proAB+*, *lacIq* ZΔM15 (TetR)] | (Atsumi, Hanai, and Liao 2008) |
| JCL260 | JCL16 Δ*ldhA* Δ*frd* Δ*adhE* Δ*pta* Δ*fnr* Δ*pflB* | (Atsumi, Hanai, and Liao 2008) |
| JCL16 Δ*lysA* | JCL16 Δ*lysA::kan* | This study |
| JCL260 Δ*lysA* | JCL260 Δ*lysA::kan* | This study |
| JCL260 Δ*lysA alsS* | JCL260 Δ*lysA::*FRT with $P_L$lac$O_1$::*alsS* integrated between *yghX* and *gpr* | This study |
| JCL260 Δ*lysA* pSA69 | JCL260 Δ*lysA::*FRT pSA69 | This study |
| JCL260 Δ*lysA alsS* Δ*galK* | JCL260 Δ*lysA alsS* Δ*galK::cat* | This study |
| JCL260 Δ*lysA yfp* | JCL260 Δ*lysA::*FRT Δ*intC::yfp-cat* | This study |
| JCL260 cm X | JCL260 Δ*lysA::*FRT with *adhA-kivD* CIChE construct integrated in the *yihG* site, maintained both on plates and in liquid medium with X µg/mL chloramphenicol | This study |
| JCL260 Δ*lysA alsS* aceK-mut | JCL260 Δ*lysA alsS mutS*- with AceK mutation P510S | This study |
| K12 Δ*ilvD* | K12 Δ*ilvD::kan* | This study |
| K12 Δ*ilvD cfp* | K12 Δ*ilvD::*FRT Δ*galK::cfp-bla* | This study |
| K12 Δ*ilvD* pSAS31 | K12 Δ*ilvD::*FRT pSAS31 | This study |
| K12 Δ*ilvD cfp* pSAS31 | K12 Δ*ilvD::*FRT Δ*galK::cfp-bla* pSAS31 | This study |
| Δ*trpB* Keio strain | BW25113 Δ*trpB::kan* (JW1253-1) | CGSC |

| | | |
|---|---|---|
| K12 Δ*hisD* | K12 MG1655 Δ*hisD::kan* | This study |
| K12 Δ*hisD* Δ*trpR* | K12 MG1655 Δ*hisD::*FRT Δ*trpR::kan* | This study |
| K12 Δ*hisD* Δ*tnaA* | K12 MG1655 Δ*hisD::*FRT Δ*tnaA::kan* | This study |
| K12 Δ*hisD* Δ*trpR* Δ*tna* | K12 MG1655 Δ*hisD::*FRT Δ*trpR::*FRT Δ*tnaA::kan* | This study |
| Δ*hisD* Keio strain | BW25113 Δ*hisD::kan* (JW2002-1) | CGSC |
| Δ*ilvD* Keio strain | BW25113 Δ*ilvD::kan* (JW5605-1) | CGSC |
| Δ*ilvE* Keio strain | BW25113 Δ*ilvE::kan* (JW5606-1) | CGSC |
| Δ*lysA* Keio strain | BW25113 Δ*lysA::kan* (JW2806-1) | CGSC |
| Δ*ppc* Keio strain | BW25113 Δ*ppc::kan* (JW3928-1) | CGSC |
| Δ*tnaA* Keio strain | BW25113 Δ*tnaA::kan* (JW3686-7) | CGSC |
| Δ*trpB* Keio strain | BW25113 Δ*trpB::kan* (JW1253-1) | CGSC |
| Δ*trpR* Keio strain | BW25113 Δ*trpR::kan* (JW4356-2) | CGSC |
| Δ*tyrA* Keio strain | BW25113 Δ*tyrA::kan* (JW2581-1) | CGSC |
| **Plasmid** | | |
| pSA65 | ColE1 ori; AmpR; $P_L$lac$O_1$::*kivd–adhA* | (Atsumi et al. 2010) |
| pSA69 | p15A ori; KanR; $P_L$lac$O_1$::*alsS–ilvC–ilvD* | (Atsumi, Hanai, and Liao 2008) |
| pSIM5 | Red expression plasmid; chloramphenicol-resistant | (Datta, Costantino, and Court 2006) |
| pSIM6 | Red expression plasmid; ampicillin-resistant | (Datta, Costantino, and Court 2006) |
| pET-ara-mCherry | Arabinose-inducible mCherry expression plasmid; kanamycin-resistant | (Park et al. 2011) |
| pSAS31 | Constitutively expressed mNeonGreen plasmid; kanamycin-resistant | (Scholz et al. 2018) Genbank accession: MK178599 |

| | | |
|---|---|---|
| pBT1-proD-mCherry | Constitutively expressed mCherry plasmid; ampicillin-resistant | Addgene plasmid #65823 |
| pTGD | CIChE plasmid; chloramphenicol- and ampicillin-resistant | (Tyo, Ajikumar, and Stephanopoulos 2009) |
| pTGD-*adhA-kivD* | pTGD with $P_L$lac$O_1$::*kivd–adhA* cloned in between the homology regions, using enzymes PstI and MluI | This study |

**Table S2: Primers and oligos**

| **Name** | **Sequence** | **Notes** |
|---|---|---|
| p15a_for | TCTGACGCTCAAATCAGTGG | 155 bp product from the p15a origin |
| p15a_rev | AGGCGTGGAATGAGACAAAC | |
| alsS_yghX_int_for | AATTTCGAAACAATGTTTCTAGTTTAGCGATTCGCCAGCGCGTATCCCGTACCGAGCGTTCTGAACAAAT | Underlined region is homologous to genome between *yghX* and *gpr*. Highlighted regions are homologous to pSA69, upstream of *kan* (for) and downstream of *alsS* (rev) |
| alsS_yghX_int_rev | GCATAAGCACGTATTTTTGCCCAGTTTTTCGTCACTCTGTGAGCCAGACTGGTGATTCCTCGTCGACCTA | |
| alsS_int_chk_front_for | ACCTCTCCTTTCCACCGTTC | Located in *gpr* gene |
| alsS_int_chk_front_rev | TCGCCTTCTTGACGAGTTCT | Located in *kan* gene |
| alsS_int_chk_back_for | GGGGAACTCATGAAAACGAA | Located in *alsS* gene |
| alsS_int_chk_back_rev | GAGATTTTCCCGTGAGCGTA | Located in *yghX* gene |
| mutS_STOP_JM | G*C*G*G*AACTGCTGTATGCAGAAGATTTTGCTGAAATGTCGTGATGATAAGGCCGTCGCGGCCTGCGCCGTCGCCCGCTGTGGGAGTTTGAA | * indicates phosphothoriated bond<br>Red indicates the introduced mutation |
| mutS_mut_as_for | CAGAAGATTTTGCTGAAATGTCGTG | Final 3' G is homologous to the introduced mutation |
| mutS_mut_as_rev | GGGTGATTTCCAGATTACGACG | |

| | | |
|---|---|---|
| mutS_seq_for | GATATCAGTTCCGGGCGTTT | |
| mutS_seq_rev | GTTCTCGACGCCAAAACC | |
| pstI_adhA_kivD_for | CTACTGCAGAATTGTGAGCGGATAACAAT | Primers for cloning $P_L lacO_1$::*adhA-kivD* into pTGD<br><br>Highlighted regions are homologous to pSA65 plasmid |
| mluI_adhA_kivD_rev | AACTACGCGTACAACAGATAAAACGAAAGG | |
| aslB_integr_for | ATGCGTCAGCATCGCATCCGGCAAAGGCAGATCTCAGCGACGAGGCAGCAGATCAATTCG | Primers for amplifying CIChE construct from pTGD-*adhA-kivD* for integration into aslB site<br><br>Underlined regions are homologous to aslB genomic locus. Highlighted regions are homologous to pTGD plasmid |
| aslB_integr_rev | CCACCACGCGCGCAGATTAAATCTGACTAAGCCGGCGCTAGCTACGGCGTTTCACTTCTG | |
| galK_cat_for | GTTTGCGCGCAGTCAGCGATATCCATTTTCGCGAATCCGGAGTGTAAGAACGTTGATCGGCACGTAAG | Underlined regions are homologous to flanking regions of *galK*. Highlighted regions are homologous to *cat*. |
| galK_cat_rev | CGGAAGAGCTGGTGCCTGCCGTACAGCAAGCTGTCGCTGAACAATATGAATTACGCCCCGCCCTGCCA | |
| aceK_510_mut_oligo | G*G*C*G*CATAGCCAGTGGCGAAACTCTTCCGGGAAAACATCGCCCGACGAGACGCTGTACCACGGTTCGCTGGCAAGTTCGTCTTCCGGATA | |
| aceK_mut_as_for | CCGTGGTACAGCGTCTCGT | Final 3' T is homologous to the introduced mutation |
| aceK_mut_as_rev | TCTGCCTTTGAGTTGGCTTT | |
| aceK_seq_for | CGCGTCTTATCATGCCTACA | |

| | |
|---|---|
| aceK_seq_ rev | TCTGCCTTTGAGTTGGCTTT |

#### Supplementary Methods

*Gene deletions, insertions and modifications*

Oligonucleotides were ordered from Integrated DNA Technologies (Coralville, IA) and are listed in Table S2.

Gene deletions were constructed by P1 phage transduction as previously described (Miller 1992), (Baba et al. 2006). Keio strains were used as the donor strains and LB agar with 50 μg/mL kanamycin was used as the selective medium. Transductants were purified from residual P1 phage by isolation streaking on LB agar supplemented with 0.8 mM sodium citrate and 50 μg/mL kanamycin, and verified by colony PCR as described previously (Baba et al. 2006). When required, the FRT-flanked kanamycin resistance gene used for selection was removed by transformation with a temperature-conditional plasmid, pCP20, expressing FLP-recombinase from a thermoinducible promoter.

To integrate the *alsS* gene into the site between *yghX* and *gpr,* the region of pSA69 containing *kan* and PLlacO1-*alsS* was amplified using primers alsS_yghX_int_for and alsS_yghX_int_rev, which contain 50 bp overhangs that add homology to the *E. coli* genome between *yghX* and *gpr* (location selected based on (Akita, Nakashima, and Hoshino 2015)). The PCR product was digested with both DpnI and SpeI for 12 h to degrade plasmid DNA. The remaining linear construct was integrated into JCL260 Δ*lysA::*FRT harboring pSIM6 by λ-Red recombineering, following published protocols (Datta, Costantino, and Court 2006).

The *adhA* and *kivD* genes were amplified from the pSA65 plasmid using primers pstI_adhA_kivD_for and mluI_adhA_kivD_rev. The product was DpnI digested to degrade residual template plasmid and cloned into the pTGD plasmid using restriction enzymes PstI and MluI. pTGD provides a chloramphenicol resistance cassette and flanking 1 kb homology regions to enable chemically inducible chromosomal evolution (CIChE). The CIChE construct was then amplified from the resulting pTGD-*adhA-kivD* using aslB_integr_for and aslB_integr_rev, which add 40 bp regions of homology to the *aslB* locus of the *E. coli* genome. After DpnI digestion, the linear construct was integrated into NV3r1 by λ-Red recombineering using pSIM6 (Datta, Costantino, and Court 2006) with selection on LB plates with 20 μg/mL chloramphenicol. A resulting integrant (named NV3r1 cm 20) were verified by PCR and Sanger sequencing. Subsequently CIChE was performed by growing NV3r1 cm 20 to saturation and passaging into in successively higher concentrations of chloramphenicol (cells were passed 1% v/v and antibiotic concentration was doubled in each passage). To construct JCL260 cm 20 Δ*lysA* pSA69 and JCL260 cm 80 Δ*lysA* pSA69 we prepared P1 lysates from NV3r1 cm 20 and NV3r1 cm 80 and transduced them into JC260 Δ*lysA::*FRT pSA69, selecting on LB plates with 10 μg/mL tetracycline, 50 μg/mL kanamycin and the corresponding concentration of chloramphenicol (20 or 80 μg/mL). Resulting colonies were isolation streaked twice on LB agar with the same antibiotics and 0.8 mM sodium citrate.

To introduce the *aceK* gene mutation into JCL260 Δ*lysA alsS*, we first knocked out the *mutS* gene using single-stranded λ-Red recombineering using pSIM5 (Datta, Costantino, and Court 2006) and oligo mutS_STOP_JM to introduce premature stop codons. We then used single-stranded λ-Red recombineering with oligo aceK_510_mut_oligo to introduce the SNP into *aceK*.

Both *mutS* and *aceK* were verified by allele-specific PCR and Sanger sequencing by the University of Michigan Sequencing Core (Ann Arbor, MI).

To knockout the *galK* gene, the *cat* gene was amplified from NV3r1 cm 20 gDNA using primers galK_cat_for and galK_cat_rev to add homology to the *galK* locus. The resulting linear construct was integrated by λ-Red recombineering with pSIM6 (Datta, Costantino, and Court 2006) into JCL260 Δ*lysA alsS*.

For fluorescent labeling of cells, K12 Δ*ilvD::*FRT was transformed with either pET-ara-mCherry or pSAS31. Secretor strains JCL260 Δ*lysA::*FRT *alsS* and JCL260 Δ*lysA::*FRT pSA69 were P1 transduced with *intC::yfp-cat* P1 lysate in order to integrate the *yfp* gene into the *intC* locus.

*Cultivation to assess production of 2-ketoisovalerate or isobutanol*

Production assessment was performed similarly to previous studies (e.g., (Atsumi, Hanai, and Liao 2008)). Overnight cultures in LB with appropriate antibiotics were diluted 1:100 v/v into 10 mL of M9IPG with 36 g/L glucose, 100 μg/mL ampicillin, and 50 μg/mL kanamycin in a 125-mL baffled, unvented polypropylene flasks. The medium was supplemented with either 5 g/L yeast extract or 3 mM lysine. Cells were grown to early exponential phase at 37 °C, 250 rpm, followed by addition of 0.1 mM IPTG. 2-KIV production cultures were further incubated at 37 °C, 250 rpm after induction. Isobutanol production culture flasks were sealed with parafilm and incubated at 30 °C, 250 rpm following induction.

*Metabolite detection*

Isobutanol and glucose concentrations were assessed by applying filtered culture broth to a Shimadzu high-performance liquid chromatograph (HPLC; model DGU 20A3R) equipped with an autosampler, Phenomenex Rezex ROA Organic Acid H+ (8%) guard and analytical columns (mobile phase, 5 mM $H_2SO_4$; flow rate, 0.6 ml/min; column temperature, 60 °C), and a refractive index detector.

2-KIV concentrations were determined using high performance liquid chromatography-quadrupole time-of-flight mass spectrometry (HPLC-Q-TOF/MS), based on the method of (Zhang et al. 2018). 5 µL filtered culture broth (diluted as needed) was applied to an Agilent 1290 Infinity HPLC system with an Agilent ZORBAX Eclipse Plus $C_{18}$ (flow rate, 0.3 mL/min, column temperature, 30 °C) column, coupled to a G6520B quadrupole-time of flight mass spectrometer (Q-TOF/MS). The mobile phase was composed of water with 0.1% formic acid (Solvent A) and acetonitrile (Solvent B). A linear gradient of 5-30% B was applied over 0-3 min, increasing to 90% B at 3.5 min. B was then maintained at 90% for 3 min, followed by a 3 min re-equilibration at 5% B. The Q-TOF/MS was operated in an electrospray ionization negative mode with gas temperature of 325 $^oC$, flow rate of 5 L/min, and nebulizer pressure of 30 psig. The capillary and fragmentor voltages were 3500 and 100 V, respectively. Reference compounds (*m/z* 112.985587 and 1033.988109) were continuously introduced into the electrospray ionization source for internal calibration. The area of the peak corresponding to 2-KIV (*m/z* 115.0401) was compared to those of standards made of known concentration of 2-KIV (Sigma-Aldrich).

*NTG mutagenesis and mutant screening*

NTG mutagenesis was performed according to the method of (Connor, Cann, and Liao 2010). The parental strain (JCL260 Δ*lysA alsS*) was cultured overnight in LB medium and then diluted 1% (*v*/*v*) into 5 mL of fresh LB and grown at 37 °C to an $OD_{600}$ of 0.5. Cells were pelleted by centrifugation at room temperature (5,000 rpm, 10 min) and washed twice with an equal volume 0.1 M Na citrate (pH 5.5) before resuspension in 0.1 M Na citrate (half the original volume). NTG was added to a final concentration of 50 μg/mL from a 1 mg/mL stock in 0.1 M Na citrate and incubated at 37 °C for 15 min. For the control experiment, an equal volume of 0.1 M Na citrate was added instead of NTG. After incubation, cells were washed twice with the original volume of 0.1 M phosphate buffer (pH 7.1). The cells were then resuspended in 5 mL of LB and incubated at 37 °C overnight for outgrowth. Before outgrowth, a small volume of cells were diluted and plated for both the NTG-treated tube and the control tube on LB plates to determine the kill count for the experiment before outgrowth. After outgrowth, cells were prepared for the agar plate screening as described in *Cell preparation for cross-feeding screening*, and plated with sensor strain on 24.5 cm square plates containing 0.5, 0.75 or 1.0 g/L norvaline, and incubated at 37 °C for 7 days. Cells were also plated on LB plates to determine LB-CFU. A volume of secretor cells corresponding to ~$6*10^4$ LB-CFU was plated on each square plate. The largest colonies from the cross-feeding screening plates were streaked on LB plates with tetracycline to isolate the secretor. Colonies from these plates were then re-screened in the microplate format of the co-culture screening. The most promising strains from the re-screening were transformed with pSA65 and subsequently tested for isobutanol production performance.

*Genomic sequencing*

The genomes of parental strain JCL260 Δ*lysA alsS* and mutant B1 strain were sequenced by the University of Michigan Sequencing Core (Ann Arbor, MI) by Illumina HiSeq-4000 using 0.5% of a lane for each strain. Genomic DNA of the strains was isolated using a Qiagen DNEasy Blood and Tissue kit. The libraries were prepared by the core, with 250 nt insert size and reads were 150 nt paired-end reads. SolexaQA++ (http://solexaqa.sourceforge.net/) was used to trim the reads. Reads were mapped to the reference genome (JCL260 Δ*lysA alsS*) using Bowtie 2 (http://bowtie-bio.sourceforge.net/bowtie2/), and SNPs were compared using SAMtools (http://samtools.sourceforge.net/).

**Microplate Assay: Additional Methods**

*Z-factors*

The Z-factor (Zhang, Chung, and Oldenburg 1999) was calculated according to following formula:

$$Z = 1 - \frac{3\sigma_p + 3\sigma_n}{|\mu_p - \mu_n|},$$

where $\mu$ and $\sigma$ are the mean and standard deviation of the $OD_{600}$-values of the positive ($p$) and negative ($n$) controls at a given time. We considered co-cultures containing JCL16 Δ*lysA* as the negative control and those containing JCL260 Δ*lysA* pSA69 as the positive control. Data from 8 replicates of each culture type were used in the calculation.

*Population composition determination of 2-KIV screening microplate cultures*

Upon reaching stationary phase, microplate co-cultures were serially diluted by factors of 10 in 1x M9 salts up to $10^{-6}$. 100 µL of $10^{-5}$ and $10^{-6}$ dilutions were plated on LB agar plates containing

Chromomax IPTG/X-Gal solution (Fisher). A variety of dilutions were plated on LB plates with tetracycline to reach ~100 colonies per plate. Each plate type was performed in duplicate. Blue colonies on X-gal plates were counted as sensor strain. Colonies on tetracycline plates, and white colonies one X-gal plate (if countable) were counted as secretor strain.

**Agar Plate Assay: Additional Methods**

*Colony composition determination*

Entire colonies were scooped off plate using inoculation loops and resuspended in M9 salts. The colony suspension was then serially diluted and plated in the same manner described for the microplate population composition determination.

*Determination of strain identity*

Secretor strains were isolated from mixed colonies by streaking on LB plates with tetracycline. The resulting colonies were then assayed by colony PCR with primers alsS_int_chk_front_for and alsS_int_chk_front_rev, which produce a short band if *alsS* is integrated, and with alsS_int_chk_front_for and alsS_int_chk_back_rev, which produce a short band if *alsS* is not integrated.

**Microdroplet Assay: Additional Methods**

*Microfluidic device fabrication*

Polydimethylsiloxane (PDMS) droplet generation and detection/sorting devices were fabricated using standard soft lithography methods. Briefly, an SU-8 photoresist (MicroChem Corp.) master mold was first created on a Si wafer by photolithography. For the detection/sorting device, multiple coating and exposure steps were required to construct flow channels and optical fiber grooves with different heights (50 μm and 80 μm, respectively). The wafer was silanized with vapor phase trichloro(1H,1H,2H,2H-perfluorooctyl)silane (Sigma-Aldrich) and the PDMS precursor was poured onto the master mold and cured at 65 °C overnight. The cured PDMS was peeled off the Si wafer, punched to form inlets/outlets, treated with oxygen plasma (Femto Scientific Inc.) for activation, and finally bonded to a glass slide to seal the device. For the droplet generation device, a 1.2 mm biopsy punch was used for both inlets a$ff$nd oulet. For the sorting device, a 1.2 mm punch was used for the droplet inlet, a 1.5 mm punch for the electrodes, and a 0.75 mm punch for the oil inlets and droplet outlets. The microelectrodes were created by flowing low melting Bi/In/Pb/Sn alloy (247 Solder) into the microchannels at 150 °C. The optical fiber (F-MCB-T-1FC, Newport Corp.) was manually embedded into the fiber groove. The sealed flow channel was flushed with trichloro(1H,1H,2H,2H-perfluorooctyl)silane at a concentration of 2% (v/v) in Novec HFE-7500 prior to use.

For the collection device (Fig. S10), a ~1 mm PDMS membrane containing a channel of height 100 μm and width 1 mm was punched with a 4-8 mm diameter hole and bonded to another PDMS slab. An inlet and outlet were punched and the PDMS was bonded to a glass slide.

*Droplet sorting*

During sorting, the detection/sorting device was placed on an inverted microscope (TS-100, Nikon). The droplets were excited by a diode laser with 450 nm wavelength, focused with a 10X objective lens, and the emission light was collected by the embedded optical fiber. The optical signal was filtered through a bandpass filter (CW525nm, BW25nm) and converted to electrical signal by photo-multiplier tubes (H9306-03, Hamamatsu). Sorting gate values for model libraries were selected based on comparison of the signal profiles from mono-secretor control sets of droplets. A real-time microcontroller-based circuit was used to trigger sorting AC voltage (400-700 $V_P$, 30khz) on the microelectrodes. A dielectrophoretic force induced by non-uniform electrical fields deflects droplets emitting positive signals into the collection channel of the device. No sorting voltage is triggered if the droplet contains signal below the specified threshold (i.e., does not contain a high number of fluorescent cells). The flow channel structure of the detection/sorting device was designed such that the negative signal droplets spontaneously enter the waste channel due to lower flow resistance. The lower flow resistance results from the larger width and shorter length of the channel. Device CAD file is available upon request. The positive channel was connected to the collection device via 0.01" ID diameter microbore tubing.

*Cell retrieval and identity determination*

Collected droplets were examined by microscopy in the collection device. The secretor identity was either determined directly by fluorescence, or by retrieving the droplets and plating on MacConkey agar plates to identify cell type by colony color. For determination by fluorescence, the droplets were inspected using a Nikon Eclipse Ti-S microscope to detect sensor strain green fluorescence (excitation filter 470/40, emission filter 525/50) and red fluorescence from mCherry (excitation filter 560/55, emission filter 675/67). A droplet containing cells displaying red fluorescence under these filters indicated the presence of the secretor strain JCL260 Δ*lysA* pSA69/pBT1-proD-mCherry and was therefore considered a positive droplet. For determination by colony count, the pool of collected droplets was retrieved by inverting the collection device and injecting HFE-7500 oil into the device to flow the droplets into an Eppendorf tube. The droplets were then chemically destabilized with 1*H*,1*H*,2*H*,2*H*-perfluoro-1-octanol. LB medium was added and the aqueous portion was plated on MacConkey agar plates containing tetracycline and 1% galactose. Plates were incubated for 48 h at 37 °C. Sensor cells did not grow due to the tetracycline. JCL260 Δ*lysA alsS* Δ*galK* produced white colonies and JCL260 Δ*lysA* pSA69 produced purple colonies. Purple colonies (indicating JCL260 Δ*lysA* pSA69) were restreaked on MacConkey agar plates with tetracycline and kanamycin (growth verified maintenance of the pSA69 plasmid) and on MacConkey agar plates with tetracycline and chloramphenicol (lack of growth confirmed that the colony is not mixed with JCL260 Δ*lysA alsS* Δ*galK*). Cells from randomly selected purple colonies were transformed with pSA65 and tested for isobutanol production performance.

#### Other Supplementary Materials

**Supplementary Movie 1:** Droplet reinjection into sorting device following cell growth. This video was recorded with a high-speed camera (Phantom Miro eX4) mounted on an inverted microscope (TS-100, Nikon) with a 10X objective lens.

**Supplementary Movie 2:** Droplet sorting following reinjection. This video was also recorded with a high-speed camera and 10X objective lens.

**Dataset S1:** SNP mutations in strain B1. Mutation coordinate refers to position in strain BW25113 (Grenier et al. 2014). The JCL260 strain is derived from BW25113.

#### Supplementary References

H. Akita, N. Nakashima, and T. Hoshino. **Bacterial production of isobutanol without expensive reagents.** Applied Microbiology and Biotechnology, 99 (2015): 991-99.

S. Atsumi, T. Hanai, and J.C. Liao. **Non-fermentative pathways for synthesis of branched-chain higher alcohols as biofuels.** Nature, 451 (2008): 86.

S. Atsumi, T.-Y. Wu, E.-M. Eckl, S.D. Hawkins, T. Buelter, and J.C. Liao. **Engineering the isobutanol biosynthetic pathway in *Escherichia coli* by comparison of three aldehyde reductase/alcohol dehydrogenase genes.** Applied Microbiology and Biotechnology, 85 (2010): 651-57.

T. Baba, T. Ara, M. Hasegawa, Y. Takai, Y. Okumura, M. Baba, K.A. Datsenko, M. Tomita, B.L. Wanner, and H. Mori. **Construction of Escherichia coli K-12 in-frame, single-gene knockout mutants: the Keio collection.** Molecular Systems Biology, 2 (2006)

M.R. Connor, A.F. Cann, and J.C. Liao. **3-Methyl-1-butanol production in *Escherichia coli*: random mutagenesis and two-phase fermentation.** Applied Microbiology and Biotechnology, 86 (2010): 1155-64.

S. Datta, N. Costantino, and D.L. Court. **A set of recombineering plasmids for gram-negative bacteria.** Gene, 379 (2006): 109-15.

F. Grenier, D. Matteau, V. Baby, and S. Rodrigue. **Complete genome sequence of *Escherichia coli* BW25113.** Genome Announcements, 2 (2014): e01038-14.

J.H. Miller. A short course in bacterial genetics: a laboratory manual and handbook for *Escherichia coli* and related bacteria (Cold Spring Harbor Laboratory Press: Plainview, NY).

J. Park, A. Kerner, M.A. Burns, and X.N. Lin. **Microdroplet-enabled highly parallel co-cultivation of microbial communities.** PloS One, 6 (2011): e17019.

S.A. Scholz, R. Diao, M.B. Wolfe, E.M. Fivenson, X.N. Lin, and P.L. Freddolino. **High-resolution mapping of a standardized transcriptional reporter reveals dedicated high and low transcription domains in the *Escherichia coli* chromosome.** Revision submitted to Cell Systems (2018).

K.E.J. Tyo, P.K. Ajikumar, and G. Stephanopoulos. **Stabilized gene duplication enables long-term selection-free heterologous pathway expression.** Nature Biotechnology, 27 (2009): 760-65.

J.-H. Zhang, T.D. Chung, and K.R. Oldenburg. **A simple statistical parameter for use in evaluation and validation of high throughput screening assays.** Journal of Biomolecular Screening, 4 (1999): 67-73.

Y. Zhang, B. Yin, R. Li, and P. He. **Determination of branched-chain keto acids in serum and muscles using High Performance Liquid Chromatography-Quadrupole Time-of-Flight Mass Spectrometry.** Molecules, 23 (2018): 147.
